# Chromosome-level genome assembly of the sacoglossan sea slug *Elysia timida* (Risso, 1818)

**DOI:** 10.1101/2024.06.04.597355

**Authors:** Lisa Männer, Tilman Schell, Julia Spies, Carles Galià-Camps, Damian Baranski, Alexander Ben Hamadou, Charlotte Gerheim, Kornelia Neveling, Eric J. N. Helfrich, Carola Greve

## Abstract

**Background:** Sequencing and annotating genomes of non-model organisms helps to understand genome architecture, the genetic processes underlying species traits, and how these genes have evolved in closely-related taxa, among many other biological processes. However, many metazoan groups, such as the extremely diverse molluscs, are still underrepresented in the number of sequenced and annotated genomes. Although sequencing techniques have recently improved in quality and quantity, molluscs are still neglected due to difficulties in applying standardized protocols for obtaining genomic data.

**Results:** In this study, we present the chromosome-level genome assembly and annotation of the marine sacoglossan species *Elysia timida*, known for its ability to store the chloroplasts of its food algae. In particular, by optimizing the Long-read and chromosome conformation capture library preparations, the genome assembly was performed using PacBio HiFi and Arima HiC data. The scaffold and contig N50s, at 41.8 Mb and 1.92 Mb, respectively, are 100-fold and 4-fold higher compared to other published sacoglossan genome assemblies. Structural annotation resulted in 19,904 protein-coding genes, which are more contiguous and complete compared to publicly available annotations of Sacoglossa. We detected genes encoding polyketide synthases in *E. timida*, indicating that polypropionates are produced. HPLC-MS/MS analysis confirmed the presence of a large number of polypropionates, including known and yet uncharacterised compounds.

**Conclusions:** We can show that our methodological approach helps to obtain a high-quality genome assembly even for a “difficult-to-sequence” organism, which may facilitate genome sequencing in molluscs. This will enable a better understanding of complex biological processes in molluscs, such as functional kleptoplasty in Sacoglossa, by significantly improving the quality of genome assemblies and annotations.

## Introduction

Studying genomes of species is essential to comprehend the biology of organisms (Salzberg, 2019). Third generation sequencing technologies, such as PacBio HiFi or Oxford Nanopore sequencing, have opened up the possibility of rapidly sequencing high-quality reference genomes of different organism groups at a reasonable price. However, sequencing methods and protocols are mainly developed and optimized for model organisms, especially human samples. In addition, DNA isolation for many non-model organisms is challenging. Sometimes even established sequencing methods do not work as expected, thus requiring special and adjusted handling (da Fonseca et al., 2016). Because of the existing bias towards developing methods for model species, certain taxonomic groups are still severely underrepresented in terms of genomic data and high-quality genome assemblies (Schell et al., 2017; Sigwart et al., 2021).

Molluscs represent the second-largest animal phylum consisting of approximately 200,000 species – many of them still undescribed (Wells, 1995; Groombridge et al., 2002; Chapman, 2011). The diversity in molluscs is not only reflected in their manifold appearances, but also in divergent life cycles and habitats. Furthermore, they are of great ecological, economic, and medical significance (Haszprunar & Wanninger, 2012; Wanninger & Wollesen, 2019). Considering their ecological and economical importance as well as species richness of the phylum, genomic resources of molluscs are still disproportionately low. It is worth noting that the number of molluscan reference genome assemblies on NCBI has more than doubled in recent years (Gomes-dos-Santos et al., 2020). However, more reference genomes need to be sequenced and annotated to shed light on the true genomic diversity and evolutionary characteristics of molluscs.

An extraordinary group within molluscs are the sacoglossans, also known as “solar-powered” sea slugs. Some sacoglossan species have evolved an exceptional photosynthetic association with chloroplasts and are able to store functional chloroplasts from their food algae in cells of their digestive gland, a process which is also referred to as functional kleptoplasty (Kawaguti, 1965; Trench et al., 1972, 1973). The role of the incorporated functional chloroplasts in the nutrition and metabolism of sacoglossan sea slugs is still highly controversial and debated among scientists. (Hinde & Smith, 1972; Hinde & Smith, 1975; Christa et al., 2014b; Cartaxana et al., 2017; Cruz et al., 2020). Most sacoglossans digest the kleptoplasts immediately (non-retention types) or after a few weeks (short-term retention types). However, six sacoglossan species are known to be capable of long-term chloroplast retention, in which the incorporated chloroplasts remain photosynthetically active for a period of two to ten months (Evertsen et al., 2007; Händeler et al., 2009; Wägele et al., 2010; Christa et al., 2014a; Wägele & Martin, 2014), such as the shallow-water Mediterranean species *E. timida* (Risso, 1818) (Bouchet, 1984; Thompson & Jaklin, 1988).

It still remains unknown how these sacoglossan species keep the chloroplasts active in their digestive gland cells without support from the algal nucleus, since chloroplasts need to import many proteins encoded by nuclear genes for their activity (Green, 2011). Polypropionate pyrones, produced in sacoglossans, have been suggested to be involved in the establishment and the maintenance of the association between sacoglossa and the incorporated chloroplasts by serving antioxidant and photoprotective roles (Torres et al., 2020). Therefore, to gain a better insight into the evolution and functionality of certain gene families, such as those encoding the polyketide synthases (PKS) involved in polypropionate pyrone biosynthesis, high-quality genome assemblies of Sacoglossa species are required in terms of completeness and contiguity.

In this work, we present the first annotated chromosome-level genome assembly of *E. timida*, and compare it with publicly available sacoglossan genomes. In addition, we provide a laboratory protocol that can be used to improve the sequencing of DNA from organisms whose sequencing is severely hampered by, for example, precipitated contaminants and DNA-bound metabolites that inhibit the sequencing polymerase. However, their sequencing can be improved by amplification-based protocols using currently available technologies and library kits (Bein et al., 2024) This newly generated high-quality genome assembly may serve as a reference genome for future genetic investigations on kleptoplasty in *E. timida* and as a high-quality resource for studies on sacoglossans and molluscs in general.

## Material & methods

### Sample collection and sequencing

Specimens of *E. timida* were collected in Cadaqués, Girona, Spain, in June 2019 (coordinates: 42.285173, 3.296461). Genomic DNA was extracted from a single individual using a CTAB-based method (Murray and Thompson, 1980). First, we prepared two PacBio ultra-low input libraries (including a PCR amplification step using polymerases A/B) using the SMRTbell^®^ gDNA Sample Amplification Kit and the SMRTbell^®^ Express Template Preparation Kit 2.0. Two SMRT cell sequencing runs were performed on the Sequel System IIe in CCS mode. In addition, to reduce potential PCR biases of the amplification polymerases A/B supplied by PacBio, we prepared two further libraries using KOD Xtreme™ Hot Start DNA Polymerase C (Merck), optimized for amplification of long and GC-rich DNA templates, in the PacBio ultra-low input protocol. These two PacBio ultra-low input libraries were each sequenced on a single SMRTcell using the PacBio Sequel IIe and Revio instruments, respectively. An initial attempt to sequence a PacBio standard/low-input library of these animals resulted in very poor sequencing results (Supplemental Table S4). The same was true for the attempt to sequence *E. timida* with Oxford Nanopore technology.

Chromatin conformation capture libraries were prepared using the Arima HiC Kit v01 (Arima Genomics) according to the manufacturer’s low-input protocol with a slight modification in the initial sample preparation steps. A whole specimen was first washed in seawater, then in deionised water, and lastly grounded with a pestle in a 1.5 mL tube. After preparing the specimen, we followed the manufacturer’s instructions for proximity ligation. The proximally-ligated DNA was then converted into an Arima High Coverage HiC library according to the protocol of the Swift Biosciences® Accel-NGS® 2S Plus DNA Library Kit. The fragment size distribution and concentration of the Arima High Coverage HiC library was assessed using the TapeStation 2200 (Agilent Technologies) and the Qubit Fluorometer and Qubit dsDNA HS reagents Assay kit (Thermo Fisher Scientific, Waltham, MA), respectively. The library was sequenced on the NovaSeq 6000 platform at Novogene (UK) using a 150 paired-end sequencing strategy, with an expected output of 30 Gb.

RNA was extracted from a whole individual from the same locality using TRIzol reagent (Invitrogen) according to the manufacturer’s instructions. Quality and concentration were assessed using the TapeStation 2200 (Agilent Technologies) and the Qubit Fluorometer with the RNA BR Reagents Assay Kit (Thermo Fisher Scientific, Waltham, MA). The RNA extraction was then sent to Novogene (UK) for Illumina paired-end 150 bp RNA-seq of a cDNA library (insert size: 350 bp) sequenced on a NovaSeq 6000, with an expected output of 12 Gb.

### Genome size estimation

The genome size was estimated following a flow cytometry (FCM) protocol with propidium iodide-stained nuclei described by Hare and Johnston (2012). Two fresh individuals of *E. timida* from Cadaqués were homogenized in a 1.5 mL tube with a pestle, and, as an internal reference standard, neural tissue from *Acheta domesticus* (female, 1C = 2 Gb) was chopped with a razor blade in a petri dish. Ice-cold Galbraith buffer (2 mL) was used as the suspension medium. The suspensions were filtered each through a 42-μm nylon mesh, then stained with the intercalating fluorochrome propidium iodide (PI, Thermo Fisher Scientific) (final concentration 25 µg/mL), and treated with RNase A (Sigma-Aldrich) (final concentration 250 µg/mL). The mean red PI fluorescence signal of stained nuclei was quantified using a Beckman-Coulter CytoFLEX flow cytometer with a solid-state laser emitting at 488 nm. Fluorescence intensities of at least 10,000 nuclei per measurement were recorded. We used the software CytExpert 2.3 for histogram analyses. After measuring the suspensions of *E. timida* and the internal reference standard separately, they were mixed. With this suspension mix, the total quantity of DNA per nuclei of *E. timida* was calculated as the ratio of the mean red fluorescence signal of the 2C peak of the stained nuclei of *E. timida* divided by the mean fluorescence signal of the 2C peak of the stained nuclei of the reference standard times the 1C amount of DNA in the reference standard. In total, two suspensions from two *E. timida* individuals were measured, each with four replicates that were measured on four different days to minimize possible random instrumental errors. The average of these eight measurements was calculated to estimate the genome size (1C) of *E. timida*. The value of the robust coefficient of variance (rCV), which should be about 5% or less, provides an estimate of the confidence level of the measurements.

Genome size and heterozygosity were estimated from a k-mer profile of the HiFi reads. First, count from Jellyfish 2.3.0 (Marçais & Kingsford, 2011) was run with the additional parameters “-F 4 -C -m 21 -s 1000000000 -t 96” and all HiFi reads as input. Second, a histogram was created from the resulting database with “jellyfish histo -t 96”. Third, GenomeScope 2.0 (Ranallo-Benavidez et al., 2020) in combination with R 4.3.1 was executed using the histogram as input. Additionally, *E. timida’s* genome size was also estimated from coverage distribution of mapped PacBio reads using ModEst (Pfenninger et al., 2022), as implemented in backmap 0.5 (Schell et al., 2017; https://github.com/schellt/backmap).

### Assembly strategy

Bioinformatic analyses were conducted with default parameters if not stated otherwise. HiFi reads were called using a pipeline, which is running PacBio’s tools ccs 6.4.0 (https://github.com/PacificBiosciences/ccs), actc 0.3.1 (https://github.com/PacificBiosciences/actc), samtools 1.15 (Danecek et al., 2021), and DeepConsensus 1.2.0 (Baid et al., 2023). All commands were executed as recommended in the respective guide for DeepConsensus (https://github.com/google/deepconsensus/blob/v1.2.0/docs/quick_start.md) except --all was applied instead of --min-rq=0.88 for ccs.

To remove PCR adapters and PCR duplicates, which might originate from the PCR amplification during the ultra-low library preparation, PacBio’s tools lima 2.6.0 (https://github.com/PacificBiosciences/barcoding) with options “--num-threads 67 --split-bam-named --same --ccs” and pbmarkdup 1.0.2-0 with options “--num-threads 84 --log-level INFO --log-file pbmarkdup.log --cross-library --rmdup” (https://github.com/PacificBiosciences/pbmarkdup) were applied, respectively.

We assembled the genome of *E. timida* from filtered PacBio HiFi reads of four SMRT cells using hifiasm 0.19.8 (Cheng et al., 2021). Subsequently, the primary contigs were processed. Contamination was filtered out by first running “screen genome” of FCS-GX 0.5.0 (Astashyn et al., 2024) with the corresponding database (downloaded on Dec 5th, 2023) and the NCBI taxonomy ID (154625). Second, “clean genome” was executed with the action report created by “screen genome” and a minimum sequence length of 1 (--min-seq-len 1). Subsequently, the FCS filtered assembly was polished using a workflow which includes DeepVariant. First, the HiFi reads used for assembly were mapped against the contigs with minimap2 2.26 (Li, 2018; 2021) and the options “-a-x map-hifi”. The bam file was sorted by coordinate with samtools 1.19.1 and duplicated HiFi reads were removed with Picard 3.1.0 (“Picard Toolkit”, 2019) MarkDuplicates and the option -- REMOVE_DUPLICATES. The assembly fasta and filtered bam files were indexed with samtools faidx and index commands, respectively. To call SNPs, DeepVariant 1.5.0 (Poplin et al., 2018) was applied. To keep only homozygous variants, SNPs were subsequently filtered using bcftools view 1.13 (Danecek et al., 2021) with the options -f ‘PASS’ -i ‘GT=“1/1”’. Then, the vcf file containing the homozygous SNPs was indexed with tabix from htslib 1.17 (Bonfield et al., 2021) to finally apply the variants inthe filtered assembly with bcftools consensus, which is from here on referred to as polished assembly. Haplotigs were purged from the polished assembly with purge_dups 1.2.6 (https://github.com/dfguan/purge_dups) together with minimap 2.24 for mapping HiFi reads and self alignment of the assembly according to the guidelines (https://github.com/dfguan/purge_dups/tree/v1.2.6?tab=readme-ov-file#--pipeline-guide), except “-x map-hifi” was applied during HiFi mapping and high coverage contigs were kept when running get_seqs (-c). Prior to HiC scaffolding, blobtoolkit 4.1.4 (Laetsch & Blaxter, 2017) was used to evaluate if contamination was still present. Taxonomic assignment for blobtools was conducted with blastn 2.15.0+ (Camacho et al., 2009) and the options “-task megablast -outfmt ‘6 qseqid staxids bitscore std’ -num_threads 96 -evalue 1e-25”. Information on coverage per contig was obtained from mapping HiFi reads used for assembly back to the assembly itself via backmap 0.5 (Schell et al., 2017; Pfenninger et al., 2022) in combination with minimap 2.26 (Li, 2018), samtools 1.17, Qualimap 2.3 (bamqc; Okonechnikov et al., 2016), bedtools 2.30.0 (Quinlan & Hall, 2010) and R 4.0.3 (R Core Team, 2020). Contigs were excluded if taxonomic assignment was not one of no-hit, Mollusca, Chordata or Arthropoda, GC was lower than 0.287, or average coverage was lower than 15.

The polished contigs that were filtered via blobtools, were scaffolded with Arima HiC reads in yahs 1.1 (Zhou et al., 2023). To do so, HiC reads were first mapped with the Arima mapping pipeline (https://github.com/VGP/vgp-assembly/blob/master/pipeline/salsa/arima_mapping_pipeline.sh), in combination with bwa mem 0.7.17 (Li, 2013), samtools 1.15.1, picard 2.27.1, and java 1.8.0 (Arnold et al., 2005). Afterwards, the bam file was processed along with the contigs in yahs. A hic file was created with the tool “juicer pre” from yahs, which was loaded together with the respective assembly file into Juicebox 1.11.08 (Durand et al., 2016; Dudchenko et al., 2018) for manual curation.

Contigs of mitochondrial origin were filtered out after manual curation. To do so, MitoHiFi 3.2.1 (Uliano-Silva et al., 2023) was applied together with the available mitochondrial genome sequence of *E. timida* (KU174946.1; Rauch et al., 2017) and the scaffolds after manual curation. Subsequently, blastn 2.15.0+ was applied to find curated scaffolds similar to our *E. timida* mitochondrial genome sequence. All scaffolds with an alignment length equal to their mitogenome length were filtered out. Additionally, all contigs flagged as circular by hifiasm were filtered out, which only included sequences that remained un-scaffolded.

During different stages of the assembly, quality controls were conducted. Basic contiguity statistics were calculated with Quast 5.2.0 (Mikheenko et al., 2018). Single copy orthologs of the provided metazoan set were searched with BUSCO 5.5.0 (Manni et al., 2021). Completeness regarding k-mers and QV values were obtained with Meryl 1.3 and Merqury 1.3 (Rhie et al., 2020). Mapping coverage distribution of HiFi reads was checked using backmap 0.5.

### Annotation

Masking of repetitive regions was conducted by identifying repeat families with RepeatModeler 2.0.5 (Bourque et al., 2018; Flynn et al., 2020) and the dependencies rmblast 2.14.1+ as search engine, and TRF 4.09 (Benson, 1999), RECON 1.08 (Bao & Eddy, 2002), RepeatScout 1.0.6 (Price et al., 2005), as well as RepeatMasker 4.1.6 (Smit et al., 2013-2015). LTR structural discovery pipeline was enabled (-LTRStruct) with the dependencies GenomeTools 1.6.2 (Gremme et al., 2013), LTR_Retriever 2.9.9 (Ou & Jiang, 2018), Ninja 0.97 (Al-Ghalith et al., 2016), MAFFT 7.520 (Katoh & Standley, 2013), and CD-HIT 4.8.1 (Huang et al., 2010). The resulting repeat families were used as repeat library in RepeatMasker 4.1.6 (Smit et al., 2013-2015) together with the options “-xsmall -no_is -e ncbi -pa 10 -s” and the dependencies rmblast 2.14.1+ as search engine, HMMer 3.4 (hmmer.org) andTRF 4.09 (Benson, 1999).

Structural annotation of the *E. timida* genome assembly, was conducted with BRAKER 3.0.8 (Stanke et al., 2006, 2008; Gotoh, 2008; Iwata & Gotoh, 2012; Buchfink et al., 2015; Hoff et al., 2016, 2019; Kovaka et al., 2019; Pertea & Pertea, 2020; Brůna et al., 2021, 2023, Simão et al., 2015; Li 2023; Huang & Li, 2023; Gabriel et al., 2021, 2023). RNAseq data from *E. timida* was mapped against the genome assembly using HISAT 2.2.1 (Kim et al., 2019), and the bam file sorted by coordinate with samtools 1.19.1 was provided to BRAKER with --bam. In addition, we downloaded the protein sets from the high-quality annotations of six molluscan species, which were then used as evidence during structural annotation: *Aplysia californica* (GCF_000002075.1; Knudsen et al., 2006), *Gigantopelta aegis* (GCF_016097555.1; Lan et al., 2021), *Mizuhopecten yessoensis* (RefSeq: GCF_002113885.1; Sato & Nagashima, 2001; Wang et al., 2017), *Octopus sinensis* (GCF_006345805.1; Yokobori et al., 2004), *Pecten maximus* (GCF_902652985.1; Kenny et al., 2020), and *Pomacea canaliculata* (RefSeq: GCF_003073045.1; Zhou et al., 2016; Liu et al., 2018) (Table 1). Before the protein sequences were concatenated and provided with --prot_seq to BRAKER, we evaluated BUSCO completeness, number of genes, and corresponding contiguity of the annotation to verify the high quality of the mollusc protein set. BRAKER was executed with the additional parameters --gff3 --threads=96 --busco_lineage=metazoa_odb10.

**Table 1.**
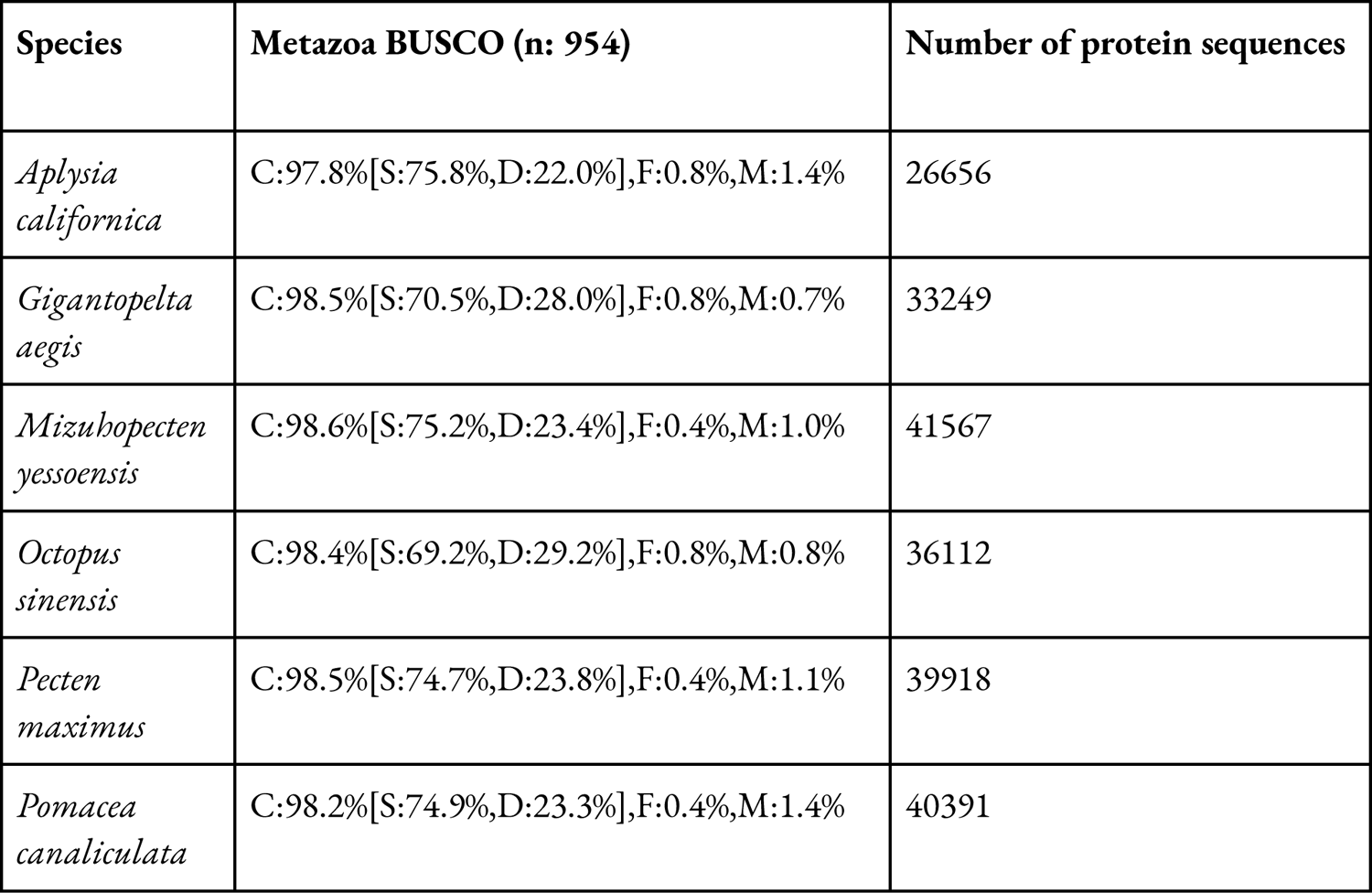
The protein sets used as evidence for the annotation of the genome assembly.

Quality controls of the *E. timida* annotation were conducted by searching for single copy orthologs with BUSCO and by calculating basic contiguity statistics of the annotated features (e.g. genes, mRNAs, etc.). The functional annotation of the predicted *E. timida* proteins was conducted using InterProScan 5.64-96.0 (Jones et al., 2014;). Databases and tools which were used during the operation with InterProScan (Paysan-Lafosse et al., 2023), are shown in detail in Supplemental Table S1. Furthermore, InterProScan was executed in the same way using the four protein sets from annotations of *Elysia chlorotica*, *Elysia crispata*, *Elysia marginata,* and *Plakobranchus ocellatus* as input (Table 4).

### Comparison of PKS encoding genes from sacoglossans

To detect the presence and copy number of PKS genes in the annotation of *E. timida,* the PKS coding sequences from the genomes of *E. chlorotica*, *Elysia diomedea* and *P. ocellatus* from Torres et al. (2020) were used as a reference (Supplemental Table S2). We also evaluated the presence and copy number of PKS genes in the annotations of *E. chlorotica*, *E. crispata*, *E. marginata* and *P. ocellatus*.

Nucleotide sequences were translated into their corresponding six protein reading frames using Geneious Prime 2023.0.4. We proceeded with the reading frames which did not contain any stop codon. These sequences were then blasted against the annotations of *E. timida*, *E. chlorotica*, *E. crispata, E. marginata*, and *P. ocellatus* using blastp 2.14.0+ (Camacho et al., 2009) and the options “-task blastp -outfmt ‘6 qseqid staxids bitscore std qlen slen’ -evalue 1e-25”. We furthermore filtered out all blast hits with an identity lower than 80%.

### Extraction and HR-HPLC-MS measurement

Individuals of *E. timida* from Cadaqués were grown in a climate chamber at 20 °C, where they spawned. Hatchling specimens (F1) were grown to adulthood in the same conditions as their parental line. The sea slugs lived in small and transparent plastic containers and under a 12:12 day-night rhythm (153 lx; λp: 545nm) (Supplemental Figure S1). Once a week water was changed and food algae (*Acetabularia acetabulum*) were supplied. Three F1 specimens were independently homogenized by blending in 0,2 Mol Tris-HCl at pH 7 in a volume of 1 mL at room temperature. The extraction was performed using ethyl acetate in tenfold excess. The crude extract was dried under reduced pressure and re-dissolved in methanol. The extract was measured with high-performance liquid chromatography-electrospray ionization high-resolution mass spectrometry (HPLC-ESI-HR-MS) using an Ultimate 3000 LC system coupled to a ImpactII QTOF (Bruker) high-resolution mass spectrometer. The extract was separated on an Acquity UPLC BEH C18 column (130 Å, 1.7 µm particle size, 2.1 mm x 100 mm) with a gradient flow of 0,4 mL/min from 5% to 95 % solvent B (acetonitrile + 0,1 % formic acid) over a time span of 14 mins. The data was acquired in positive mode at a scan range between *m/z* 100 to *m/z* 1200 and analyzed using the Bruker software DataAnalysis 4.3 and MetaboliteDetect 2.1.

### Molecular networking and visualization

The HPLC-MS/MS datasets obtained from analysis of the crude extracts from *E. timida* specimens were uploaded to Global Natural Products Molecular Networking (GNPS) to generate a molecular network, setting the minimum matched peaks to 7 and a cosine score of 0.6 (Wang et al., 2016). The software Cytoscape 3.9.1 was used to visualize the molecular network that was generated (Shannon et al., 2003). The excerpt of the network that represents the polypropionate compounds was identified by the presence of nodes representing characterized compounds (Torres et al., 2020). The compounds were identified based on the mass-to-charge ratio and sum formula. Also similar fragmentation patterns among the compounds were taken into consideration as a marker for a putative polypropionate. Only compounds with sum formulas containing exclusively carbon, oxygen and hydrogen and MS/MS fragmentation patterns indicate the presence of multiple methyl groups, clustered with the characterized polypropionates at the selected parameters.

### Construction of proposed sequence for EtPKS1 mRNA

The putative sequence of the mRNA for *E. timida* PKS1 (EtPKS1) was proposed based on sequence similarity with the transcript for the EcPKS1 (Accession number: MT348433). The conserved “GHSMGE” motif in the acyltransferase domain of PKS1 was identified on nucleotide level and used as a bait for the identification of the genomic area encoding the EtPKS1 (Torres et al., 2020). An excerpt of the genomic sequence around this sequence motif was translated in the three forward translation frames to map the translations with the EtPKS1 transcript and annotate the exons.

## Results

### Genome size estimation

Flow cytometry results were represented as histograms displaying the relative propidium iodide fluorescence intensity which we received after a simultaneous analysis of *E*. *timida* 2C and the house cricket *A. domesticus* 2C as an internal reference standard (Supplemental Figure S2). The obtained average genome sizes for the two individuals of *E. timida* were 898.00 and 891.78 Mb, respectively. From all eight measurements, including both individuals, a genome size of 894.89 Mb was estimated (Supplemental Table S3).

Mapping based genome size estimation with ModEst based on the final genome assembly resulted in 632 Mb. Based on k-mers, the genome size was estimated to be 548,2 Mb and heterozygosity 0.794% (Supplemental Figure S3). The heterozygosity value of *E. timida* is higher compared to other sacoglossan species (Supplemental Table S4).

### Sequencing

The four PacBio sequencing runs using the standard PacBio ultra-low and the modified PacBio ultra-low input library preparations yielded a total polymerase read length of 510 Gb, 431 Gb, 612 Gb, and 1360 Gb, respectively (Supplemental Table S5 and Supplemental Figure S4). Illumina sequencing of Arima HiC and RNAseq libraries resulted in 95.7 and 40.6 million read pairs, corresponding to 28.7 and 12.2 Gb, respectively.

PacBio Hifi reads and Arima HiC reads which were used for genome assembly, RNAseq reads as well as the final assembly and annotation can be publicly accessed via BioProject PRJNA1119176 and this link: https://genome.senckenberg.de/download/etim/

### Assembly

HiFi calling and subsequent PCR duplicate removal resulted in more than 19 million HiFi reads with a total length of 108 Gb and an N50 of 6143 bp (more details in Supplemental Figure S4). Given the genome size estimates based on FCM (895 Mb) and ModEst (632 Mb), the theoretical coverages were calculated to be 120x and 170x, respectively. FCS-GX identified 873 sequences (53.3 Mb) containing contamination, which were excluded (871) or trimmed (2). Nevertheless, after polishing and purging, the blobplot still showed the presence of contamination (Supplemental Figure S5). In total, 3084 additional contigs (100.8 Mb) were removed before proceeding with HiC scaffolding (blobplot after removing contamination in Supplemental Figure S9). After manual curation, scaffolding resulted in 15 chromosome-scale sequences (Figure 1), showing no contamination (Supplemental Figure S6). These 15 scaffolds made up 89.1% of the assembly’s total length. The total length of the final *E. timida* genome assembly was 754 Mb with a scaffold and contig N50 of 41.8 Mb and 1.92 Mb, respectively. Furthermore, the assembly had a QV of 58.3. Additional quality metrics of the final *E. timida* assembly and a comparison to publicly available sacoglossan genome assemblies are shown in Figure 2 and Table 2.

**Figure 1.**
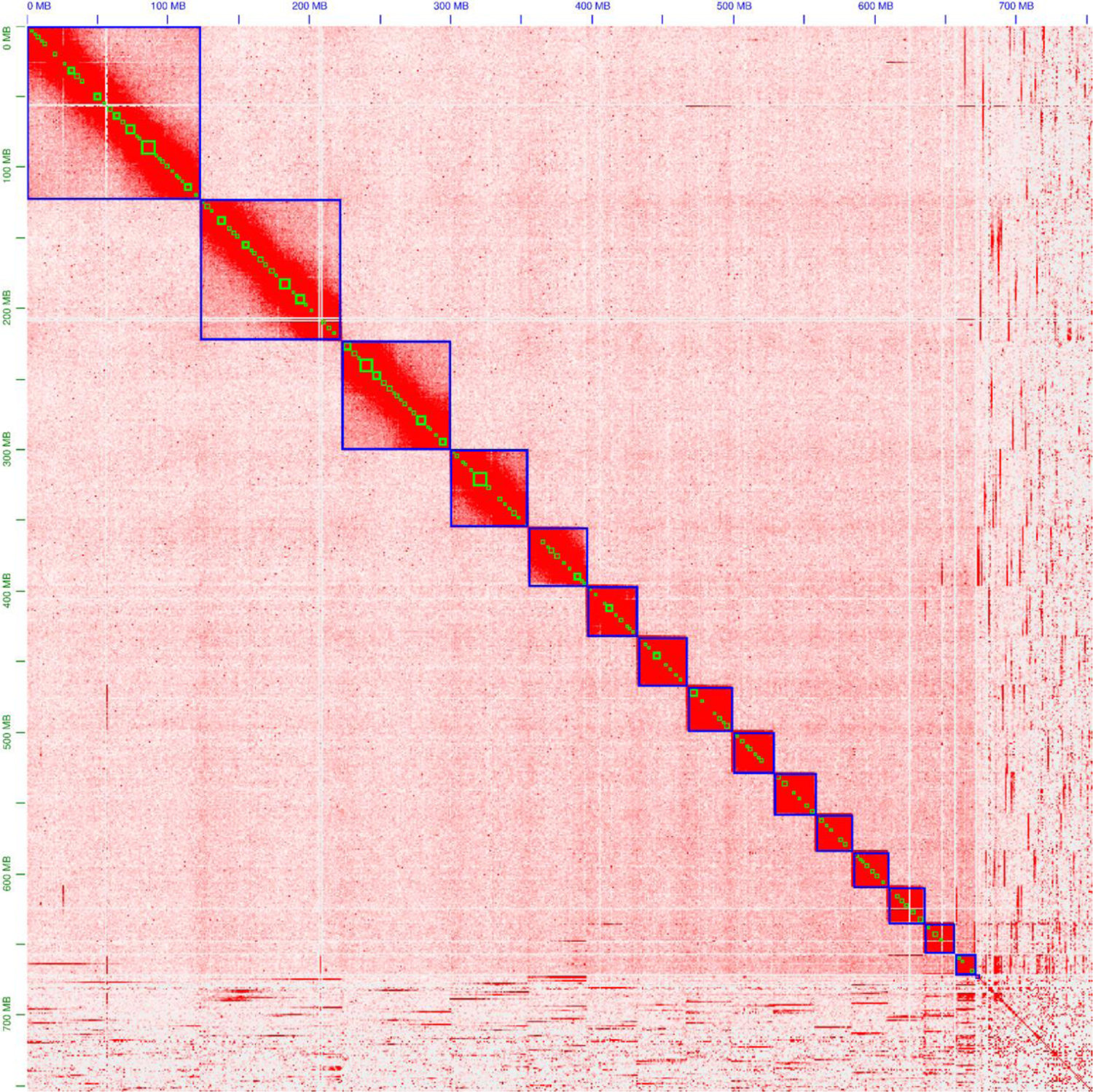
Contact map after yahs scaffolding using Arima HiC data and manual curation. Blue and green squares mark scaffolds and contigs respectively. Higher number of contacts is represented by higher intensity of the color.

**Figure 2.**
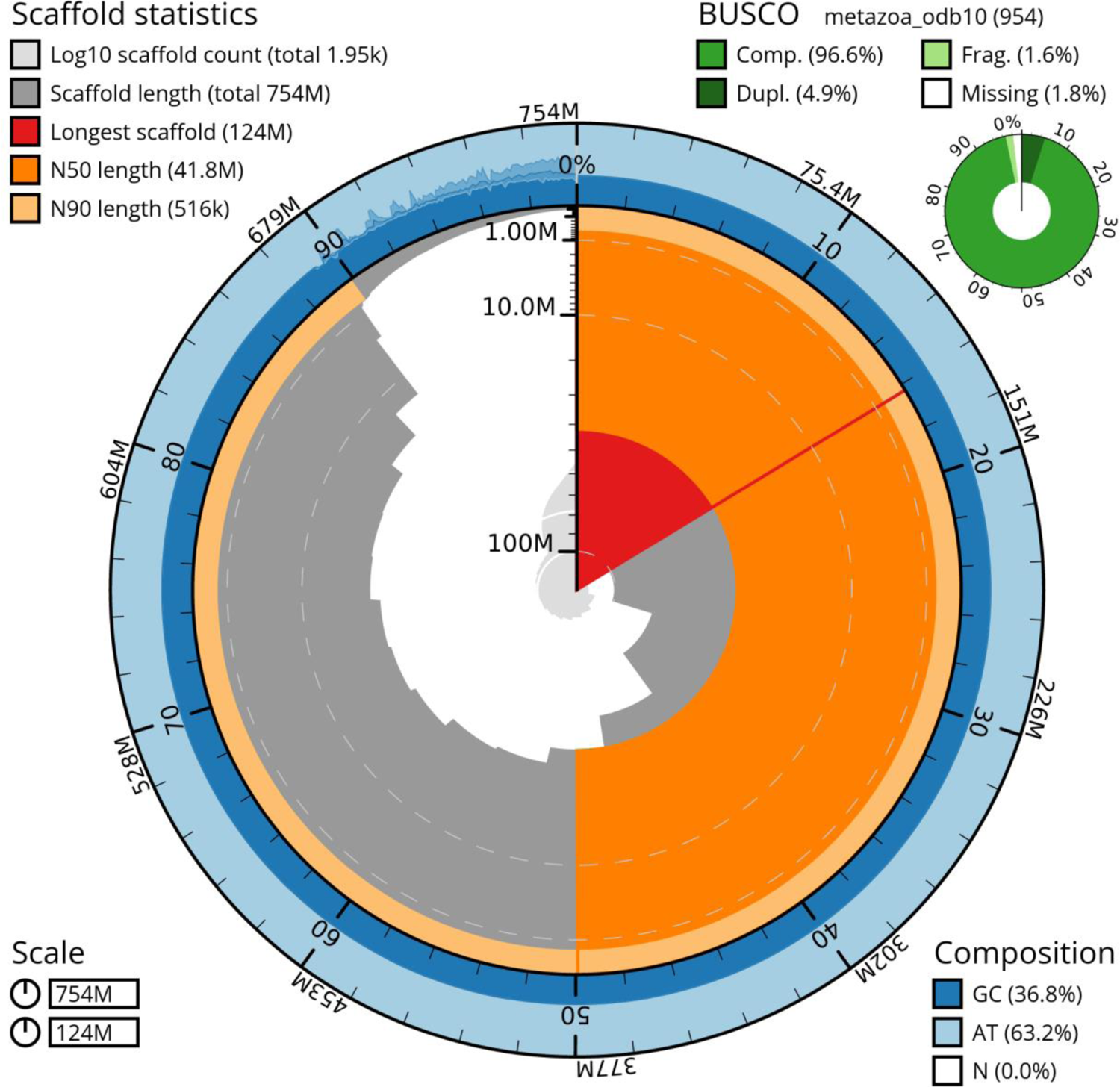
Snail plot of the final genome assembly. The plot created with blobtoolkit visualizes amongst others scaffold count, lengths, length distribution, nucleotide composition, and recovered BUSCOs.

**Table 2.**
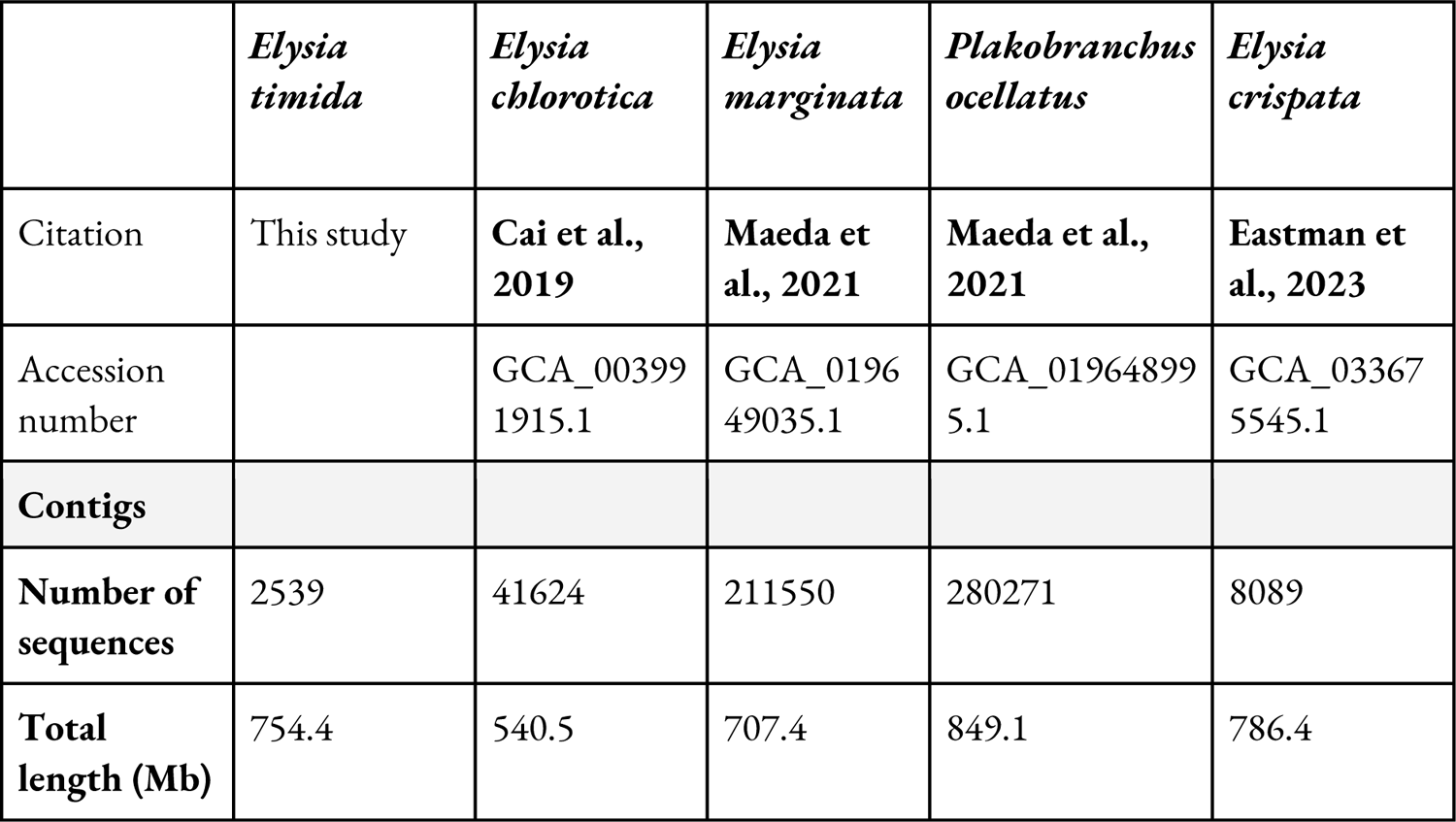
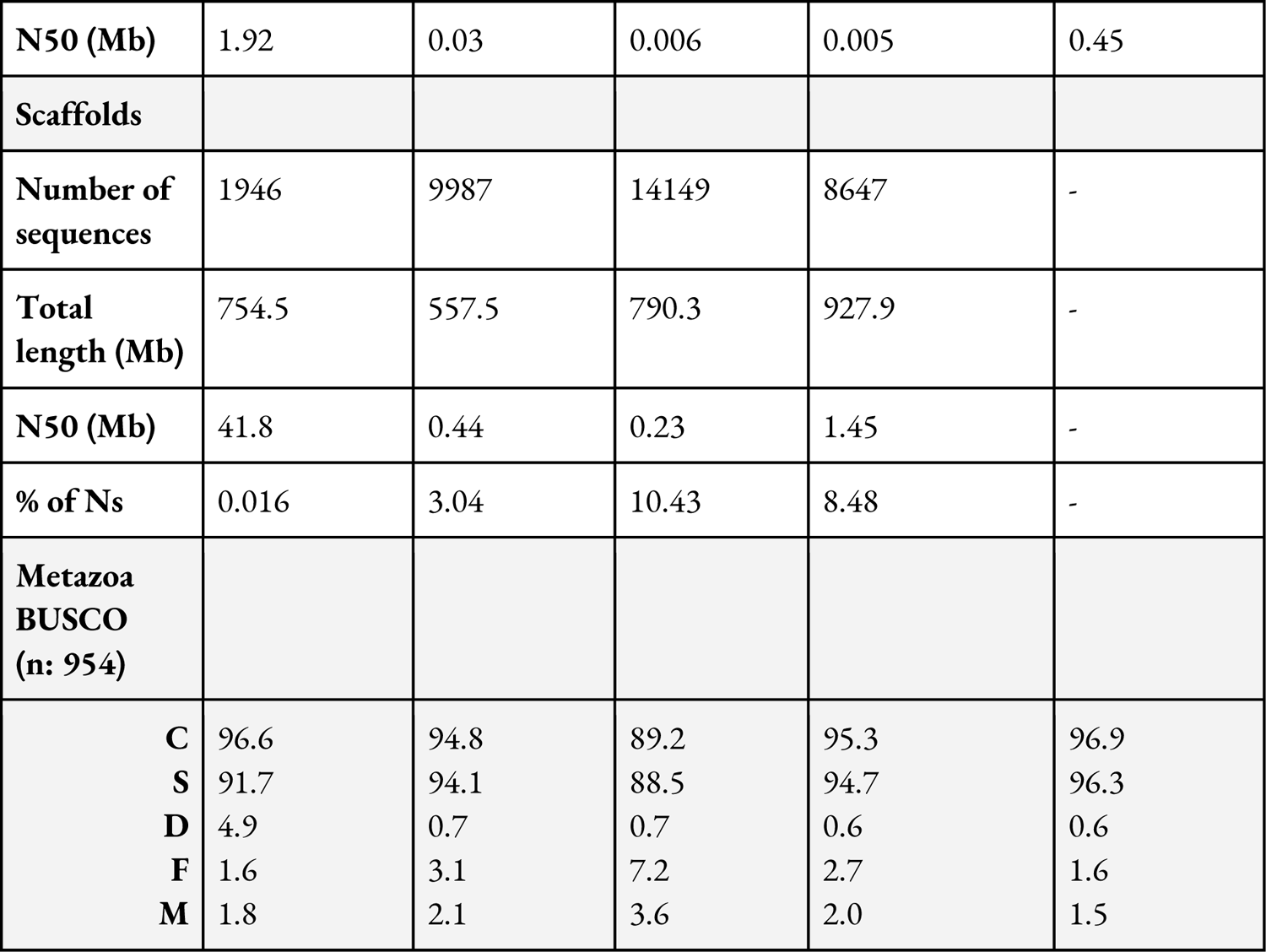
Quality metrics of the *E. timida* genome assembly in comparison to available sacoglossan species. The BUSCO values are given in %.

The contig N50 of the *E. timida* genome assembly is fourfold higher compared to the highest achieved so far sacoglossan species’ genome assembly (*E. crispata*: 0.45 Mb), while BUSCO values of *E. timida* were similar to the other sacoglossans (Table 2). Only duplicated BUSCOs were higher for *E. timida* compared to the other assemblies. As no HiC data has been generated for other Sacoglossa before, the *E. timida* assembly has a manyfold higher scaffold N50 in comparison (Table 2).

### Annotation

RepeatModeler identified sequences of 1,886 repeat families with a total length of 1,705,165 bp. Subsequently, RepeatMasker annotated 44.3% of the assembly as repetitive, of which the majority of repetitive families were labeled as unclassified (32.5% of the assembly). Contiguity statistics and BUSCO results of the *E. timida* annotation compared to annotations of other Sacoglossa are in Table 3. The values of the complete BUSCOs range between 86.1% and 92.9% which confirm the high quality of the protein sets of published sacoglossan annotations.

**Table 3.**
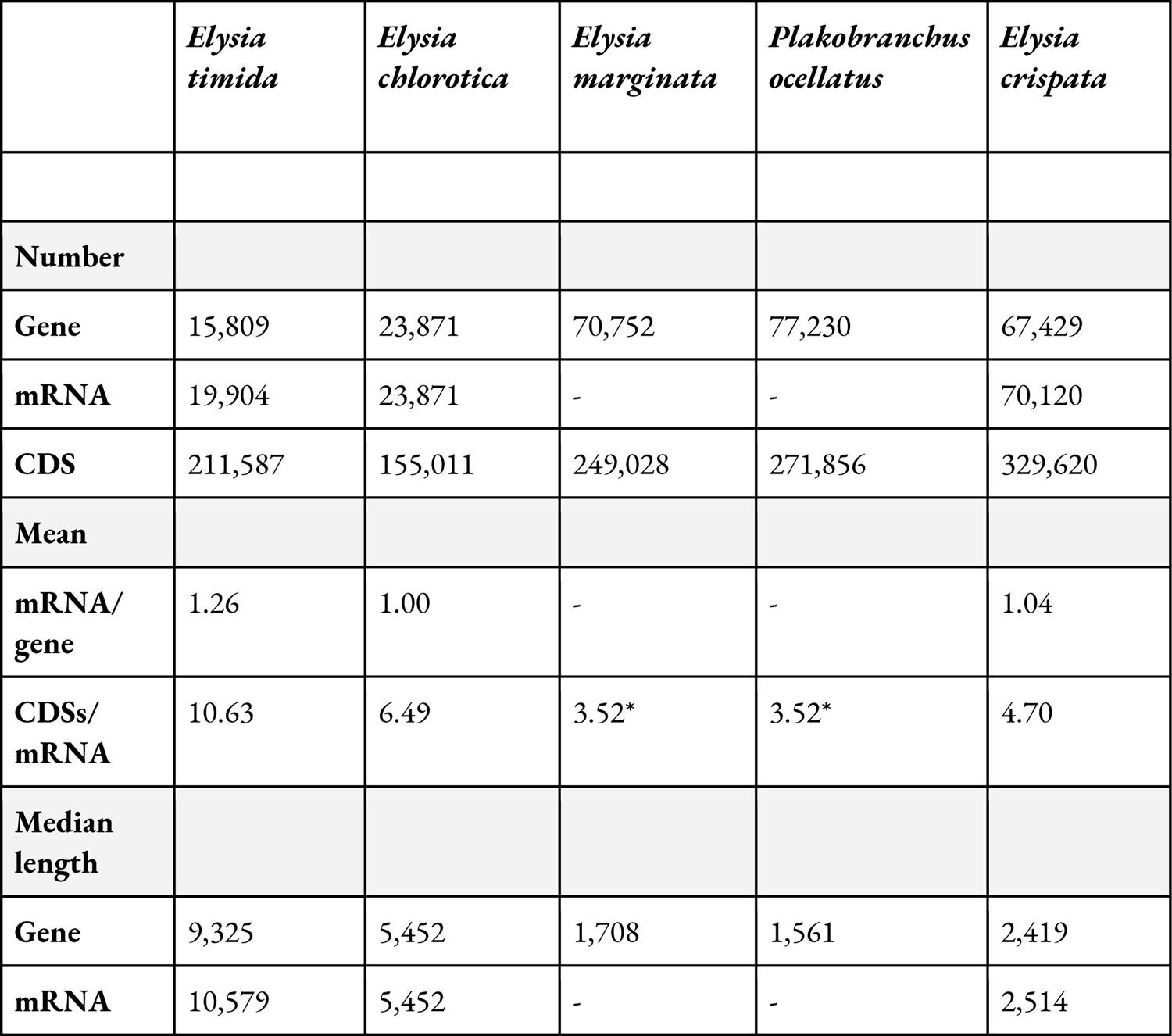
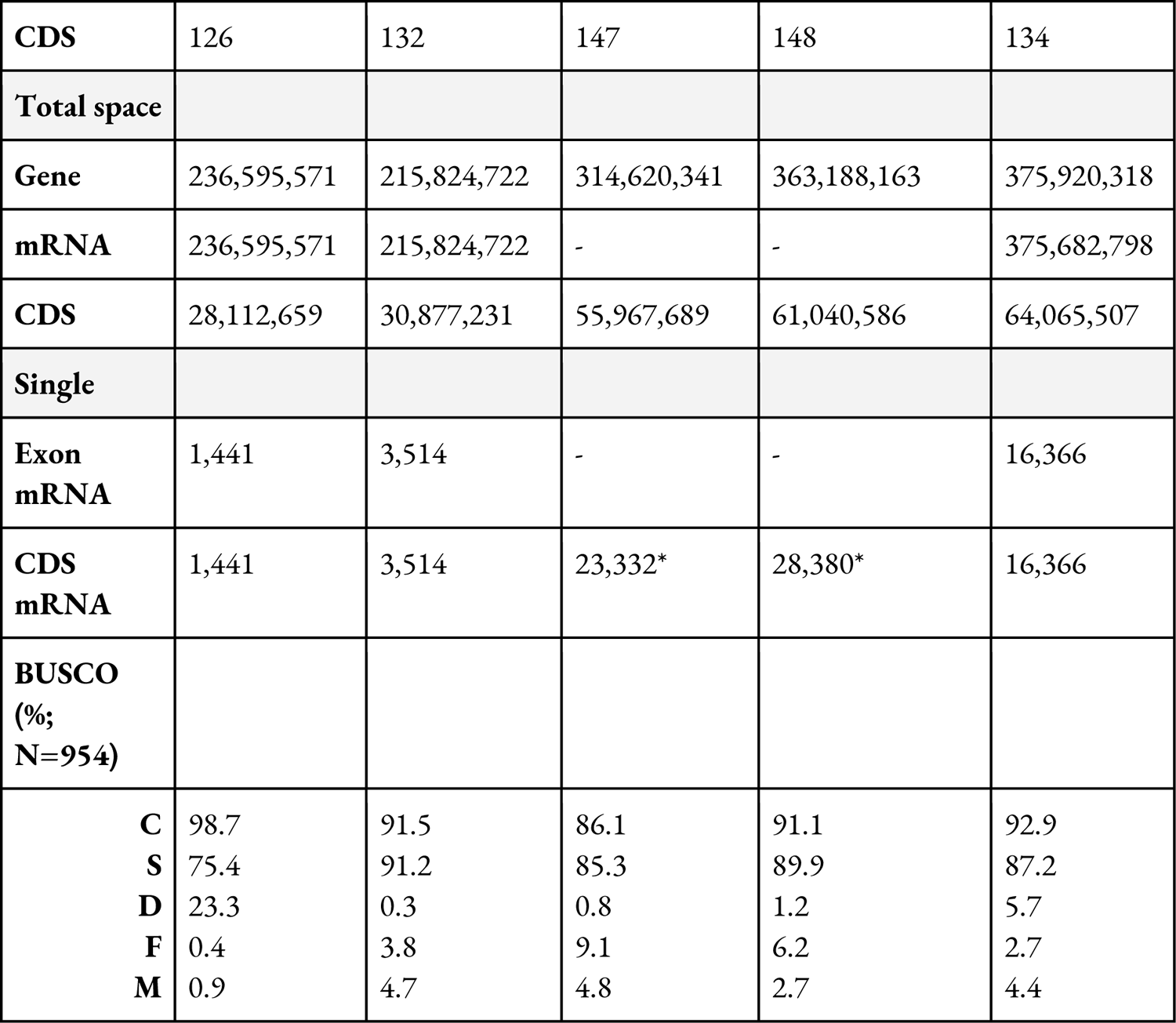
Quality metrics of the *E. timida* genome annotation in comparison to available sacoglossan species. *For Annotations of *E. marginata* and *P. ocellatus* only “gene” and “CDS” features are annotated, so that no metrics regarding mRNAs and mean CDSs/mRNA as well as single CDS genes are shown.

**Table 4.**
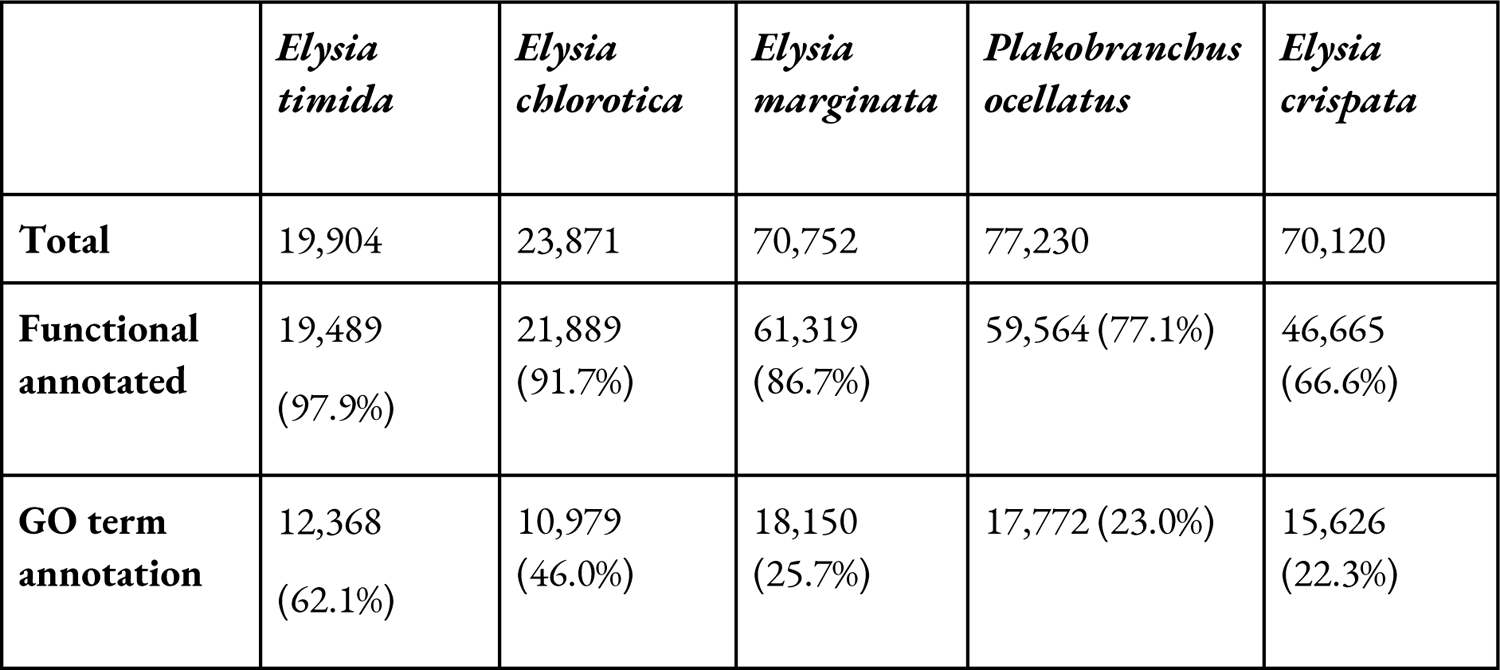
Number of protein sequences annotated structural (Total) and functional with at least one analyses in InterProScan or with at least one GO term.

In comparison to annotations of other sacoglossan genome annotations, *E. timida* was the most complete and least fragmented in terms of BUSCOs. Additionally, mean CDS per mRNA and median gene length among others was also the highest for *E. timida*, whereas the number of single CDS mRNAs was the lowest. Total gene and CDS space were similar or lower compared to other annotated sacoglossan species.

InterProScan assigned at least one functional annotation with any of the applied analyses to 19,489 (97.9%) different *E. timida* protein sequences. Furthermore, 12,368 (62.1%) different *E. timida* protein sequences were annotated with at least one Gene Ontology (GO) term, whereas this percentage was lower in other Sacoglossa with a larger range (from 22 to 46%). A comparison of functional annotated protein sequences to other Sacoglossa can be found in Table 4.

### PKS presence in *E. timida* and comparison of PKSs in sacoglossans

All FAS and PKS encoding genes from *E. chlorotica*, *E. diomedea* and *P. ocellatus* (Torres et al., 2020) were found to be annotated for the *E. timida* genome assembly before filtering. We also checked if we find all PKS encoding genes in the annotations of the sacoglossans *E. chlorotica*, *E. crispata*, *E. marginata* and *P. ocellatus*. Before filtering, the nine PKS or FAS encoding genes from three sacoglossan species were detected in all of the other sacoglossans. After we filtered out all the blast hits with an e-value above 1e-25 and a percentage of identical positions below 80%, the number of hits reduced drastically in all sacoglossans (Supplemental Table S6). Before filtering, between two and 38 gene hits were found in all sacoglossan species. However, after filtering, the maximum of gene copies was found in the annotation of *E. marginata* in which we had four blast hits of the EcPKS1 gene. Some sacoglossan annotations did not have any FAS or PKS gene hits after filtering. The EtFAS, EtPKS1 and EtPKS2 encoding genes were annotated in the genome of *E. timida* (Table 5). The genes encoding the EtFAS and EtPKS2 were annotated automatically by BRAKER 3.0.8. In contrast, the gene encoding the EtPKS1 was not recognized automatically, and was annotated manually based on the alignment with the EcPKS1 amino acid sequence (Supplemental Figure S7). The presence of PKS encoding genes in the genome of *E. timida* was investigated to get an indication of which polypropionates can be produced by the sea slug.

**Table 5.** Sequence identifiers for the nucleotide and amino acid sequences of EtFAS, EtPKS1 and EtPKS2.

| **Coding region in genome assembly** | **Transcript level** | **Protein level** |
| --- | --- | --- |
| Scaffold_4:11989323-12013722 | g10811.t1 | EtFAS |
| Scaffold_6:1080882-1152789 | Assembled manually | EtPKS1 |
| Scaffold_12:1113801-1144318 | g4188.t1 | EtPKS2 |

### Identification of putative polypropionates

To investigate the spectrum of polypropionates produced by the sea slug, three adult specimens were extracted. Analysis of HPLC-MS/MS data of the crude resulted in the identification of putative polypropionates (Supplemental Figure S8). The characteristic MS/MS fingerprints of characterized polypropionates were used to identify related compounds based on their similar MS/MS fragmentation patterns. The obtained HPLC-ESI-MS data was visualized to represent the relatedness between the putative compounds (Figure 3). Each node represents a polypropionate and is labelled with the detected mass-to-charge ratio. The edge lines visualize the relatedness between two compounds indicating that they have similar structures. More than half of the 18 detected putative polypropionates matched characterized compounds that were reported from different *Elysia* and *Plakobranchus* species (Table 6) (Li et al., 2023). Additionally, eight putative polypropionates that have not been structurally characterized were detected.

**Figure 3.**
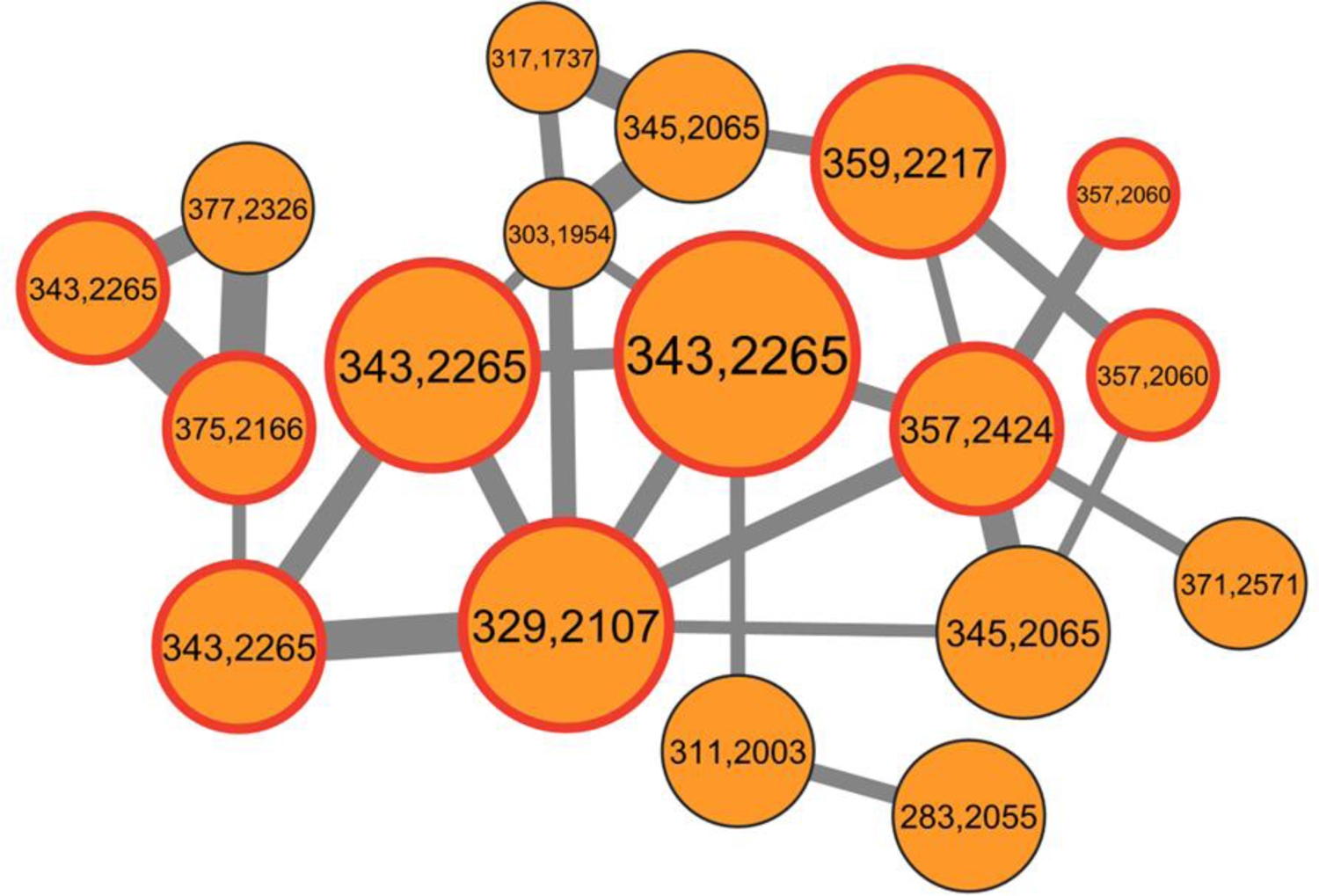
Excerpt of a molecular network showing detected polypropionates from crude extracts of *E. timida*. Putative polypropionates were clustered based on their similar MS/MS fragmentation patterns. Each node represents a polypropionate and is labeled with the detected mass. Masses that correspond to characterized polypropionates are highlighted in red. The node size corresponds to the production level and the edge width represents the relatedness between two compounds. The thresholds for the cluster were set to 7 minimum matched peaks and a cosine score of 0.6.

**Table 6.**
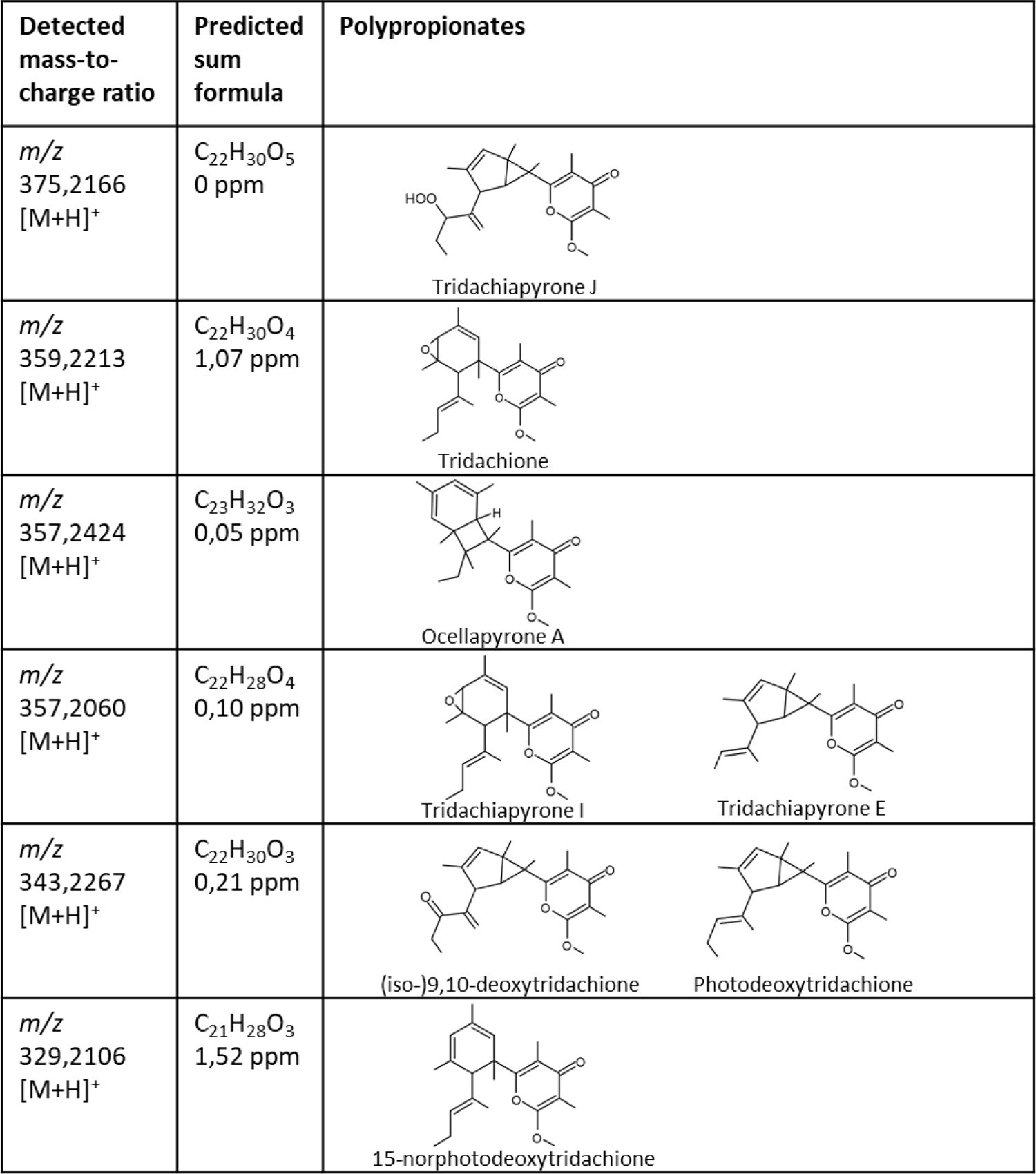
Overview of polypropionates isolated from kleptoplastic sacoglossans. Compounds were detected by LC-HR-MS/MS analysis of crude extracts. Only masses corresponding to characterized polypropionates are shown, new putative compounds were excluded.

| Detected mass-to-charge ratio | Predicted sum formula | Polypropionates |  |
| --- | --- | --- | --- |
| $m/z$<br>375,2166<br>[M+H]<sup>+</sup> | C<sub>22</sub>H<sub>30</sub>O<sub>5</sub><br>0 ppm | Tridachiapyrone J | |
| $m/z$<br>359,2213<br>[M+H]<sup>+</sup> | C<sub>22</sub>H<sub>30</sub>O<sub>4</sub><br>1,07 ppm | Tridachione | |
| $m/z$<br>357,2424<br>[M+H]<sup>+</sup> | C<sub>23</sub>H<sub>32</sub>O<sub>3</sub><br>0,05 ppm | Ocellapyrone A | |
| $m/z$<br>357,2060<br>[M+H]<sup>+</sup> | C<sub>22</sub>H<sub>28</sub>O<sub>4</sub><br>0,10 ppm | Tridachiapyrone I | Tridachiapyrone E |
| $m/z$<br>343,2267<br>[M+H]<sup>+</sup> | C<sub>22</sub>H<sub>30</sub>O<sub>3</sub><br>0,21 ppm | (iso-)9,10-deoxytridachione | Photodeoxytridachione |
| $m/z$<br>329,2106<br>[M+H]<sup>+</sup> | C<sub>21</sub>H<sub>28</sub>O<sub>3</sub><br>1,52 ppm | 15-norphotodeoxytridachione | |

Many of the characterized compounds harbor stereocenters which were not considered for the assignment of the putative polypropionates to known structures as they cannot be determined by mass spectrometry. After production of the polypropionates by the PKS, tailoring reactions such as cyclizations and epoxidations take place. This results in the wide spectrum of polypropionates produced by sacoglossans (Supplemental Figures S7 and S9) (Li et al., 2023). The genes encoding tailoring enzymes were not clustered with the PKS encoding genes.

## Discussion

The adaptive potential and remarkable survival mechanisms of Sacoglossa are subject of many studies. The inclusion of fully annotated Sacoglossa genomes in these studies is essential to properly investigate the genetic processes underlying functional kleptoplasty and to understand its functional role. Our *E. timida* genome assembly achieves the highest values of contiguity, of BUSCOs completeness and accuracy compared to available sacoglossan genome assemblies and therefore makes a valuable contribution to this understanding. In addition, we identified genes encoding PKS1 and PKS2 in the genome annotation of *E. timida*, suggesting that *E. timida* is a putative producer of polypropionates. It is hypothesized that polypropionates are involved in the establishment and maintenance of the association between Sacoglossa and the incorporated chloroplasts (Torres et al., 2020). Ireland and Scheuer (1979) found out that in Sacoglossa fixed carbon which was acquired from the photosynthesis of *de novo* chloroplasts, was embodied into polypropionates. The polypropionates act via oxidative and photocyclization pathways which are suggested to behave like sunscreen and prevent the sea slugs from photosynthetic damage (Ireland and Scheuer, 1979; Zuidema et al., 2005; Zuidema & Jones, 2005; Zuidema & Jones, 2006; de Vries et al., 2015; Powell et al., 2018). Polypropionates would therefore be needed in species with a photosynthetic lifestyle, such as *E. timida*.

Despite their high diversity, molluscs are still very poorly studied in terms of publicly accessible high-quality reference genomes, partly due to the aforementioned difficulties in DNA extraction, library preparation and sequencing (van Bruggen et al., 1995, Stork, 1999, Ponder & Lindberg, 2008). Currently available molluscan genome assemblies in the National Center for Biotechnology Information (NCBI) cover only ∼0.1% of the described species in the entire phylum. One reason for the often rather fragmented genome assemblies in molluscs could be that the sequencing polymerases are hindered by contaminants, such as the polysaccharide-containing mucus of molluscs, or metabolites bounded to the DNA (Aoki & Koshihara, 1972; Sokolov, 2000). The sacoglossan genome assemblies published so far are quite fragmented, with contig N50s of 0.005 to 0.45 Mb (Cai et al., 2019, Maeda et al., 2021, Eastman et al., 2023). The reason is that in most cases long-read sequencing, but also the chromatin conformation capture library preparation did not work, as so often in molluscs. This resulted in very low sequencing yield. To enable the sequencing of these animals, we have therefore switched to the PacBio ultra-low input protocol, which includes a PCR amplification step to increase the amount of DNA relative to possible contaminants and to obtain ‘artificial but clean’ DNA that can then be easily sequenced. In general, PCR amplification used in the preparation of ultra-low input libraries can lead to bias towards some genomic regions. However, using different PCR polymerases for amplification can counteract this bias and complementary amplify different genomic regions. Combining these data thus leads to improved contiguity of genome assemblies (Bein et al., 2024). By using the Arima HiC low-input library preparation protocol, higher cross-linking yields were achieved, ultimately resulting in higher coverage and improved insert sizes (compared to OmniC (data not shown)). With a scaffold and contig N50 of 41.8 Mb and 1.92 Mb respectively, the *E. timida* genome assembly of this study is 100-fold and 4-fold higher than the other sacoglossan genome assemblies, respectively.

However, there is a discrepancy of total assembly length (754 Mb) and genome size estimates (FCM: 895 Mb; ModEst: 632 Mb; GenomeScope: 548 Mb). The smaller total length compared to the FCM estimate may be due to collapsed repeats that have not yet been resolved. Although PacBio sequencing was successful, the overall N50 of the HiFi reads (6 kb) may, in some cases, still be too short to resolve some long repetitive regions of the genome. The mapping-based genome size estimate of ModEst is considerably smaller than the FCM estimate and the total assembly length. ModEst assumes that differences in coverage are due to technical problems in assembly. This may not be entirely correct in our case, as differences in coverage may be caused by bias in PCR amplification. Due to its assumption, the ModEst estimate appears to be less reliable than the FCM estimate with rCV values < 5 %. Similarly, genome sizes appear to be consistently underestimated by k-mer-based methods, partly due to repeats (Pfenninger et al., 2022). Therefore, a comparison between the total length of the high-complexity regions in the assembly and a k-mer-based estimate of genome size is useful. Adding the number of masked bases in the assembly (335Mb) and the k-mer-based genome size estimate (548Mb) yields 883Mb, which is very close to the FCM-based genome size estimate (895Mb). The number of chromosome-level sequences of 15, obtained from the HiC data for *E. timida*, is consistent with the karyotype of the sacoglossan species *Oxynoe olivacea* (Vitturi et al., 2020). In earlier publications, 17 chromosomes were found in other more closely related sacoglossan species (Mancino & Sordi, 1965; Burch & Natarajan, 1967). However, the karyotype of *E. timida* was not included and needs to be verified in future studies.

The BUSCO values were similar to other sequenced sacoglossan species. However, the duplicated BUSCOs in the *E. timida* genome assembly were higher (4.9 %) than in the other genome assemblies (<1 %). These duplications can be caused by true biological events, replicating loci which contain genes thought to be single copy orthologs. In addition, duplicated BUSCOs can result from high heterozygosity, resulting in the same genomic locus (in a diploid organism) being assembled twice, creating so-called haplotigs. Haplotypic duplications are searched for and collapsed with both hifiasm and purge_dups (here only at the ends of contigs). The heterozygosity of the *E. timida* genome (0.794%) is estimated to be higher than in other Sacoglossa (0.18%-0.42%; Supplementary Table S6; Theisen & Jensen, 1991). However, with only 4.9% duplicated BUSCOs, the proportion is still quite low.

Contamination can lead to major problems when dealing with genome assemblies from public databases (Astashyn et al., 2024; Steinegger & Salzberg, 2020; Challis et al., 2020). Therefore, contamination screening is a fundamental part of the genome assembly. The presented assembly of *E. timida* was screened with two different contamination tools. The advantage of blobtools over the sequence similarity based method FCS-GX was that coverage and GC content are taken into account additionally. In cases where taxa are underrepresented in a database for sequence similarity searches (e.g. Mollusca), false positive and false negative hits occur more frequently. In addition, shorter sequences are less likely to be identified in general. However, taking into account the read coverage and GC content, short sequences can still be identified as contaminants with a high probability (see cluster at the bottom left of Supplemental Figure S10). False-positive hits are still possible due to the taxonomic assignment by blobtools. Nevertheless, we did not filter out sequences that were assigned to Chordata or Arthropoda because the nt database does not contain the necessary diversity of Mollusca sequences to reliably identify this phylum in a *de novo* assembly. The contigs of the *E. timida* assembly therefore generate hits for the closest related species in the database, likely due to conserved elements of the genome across different phyla (e.g. protein domains).

The structural annotation of the *E. timida* genome assembly shows excellent quality metrics and is the most complete in terms of BUSCOs compared to available annotated genome assemblies of Sacoglossa. In particular, the higher number of CDSs/mRNA and the longer median gene length, while total gene space is comparable, indicate a higher contiguity of annotated genes. This seems to be strongly influenced by the contiguity and accuracy of the underlying genome assembly. Furthermore, over 60% of the protein sequences resulting from the *E. timida* annotation were annotated with at least one GO term, which is 1.4- to 2.8-fold higher compared to the annotation of other Sacoglossa (Table 4). The absolute numbers of protein sequences annotated with a GO term are probably higher for other Sacoglossa due to gene fragmentation. When a gene is split between two contigs or scaffolds, it is annotated as two different genes, but if both parts are large enough to make a reliable match, a GO term (maybe even the same one) is assigned to both gene fragments. The overall functional annotation rate is lower for the other Sacoglossa, probably due to general fragmentation and sequences becoming too short to reliably match against protein sequences of known functions. However, the high number of genes annotated in other Sacoglossa may not be due to fragmentation of the assembly alone. To show that many genes could be true or false positive annotations, orthologous clustering could be performed with all available Sacoglossa and other high-quality annotations from other molluscs. As many downstream analyses (e.g. comparative, evolutionary) depend on high-quality data as input, the presented annotation will enhance or even enable future studies.

Although it has been claimed that polypropionates are not produced by the animals themselves, but by symbiotic bacteria or dietary organisms (Morita & Schmidt, 2018; Zan et al., 2019), there is increasing evidence that animals are capable of producing various compounds themselves (Castoe et al., 2007; Shou et al., 2016; Cooke et al., 2017; Sabatini et al., 2018; Torres & Schmidt, 2019). In Sacoglossa, it is assumed that the produced polypropionate pyrones contribute, among other things, to the establishment and maintenance of the association of Sacoglossa and incorporated chloroplasts (Torres et al., 2020). The identification of genes encoding PKS1 and PKS2 in the genome annotation of *E. timida* indicates that *E. timida* is a likely producer of polypropionates. This makes sense considering that polypropionates are thought to act like a sunscreen and protect *E. timida* from photosynthetic damage (Ireland and Scheuer, 1979; Zuidema et al., 2005; Zuidema & Jones, 2005; Zuidema & Jones, 2006; de Vries et al., 2015; Powell et al., 2018). Depending on the genome annotation and the origin of the proteins, FAS, PKS1 and PKS2 genes were found in the sacoglossan assemblies. The quality of the genome assemblies and the protein sequences seem to have a major influence on the number of gene copies found. No gene copies of the protein sequences of *P. ocellatus* were found in the annotation of *E. timida*, which could be due to the fact that *P. ocellatus* is a more distant related species to *E. timida* than the other *Elysia* species. The gene sequences encoding FAS, PKS1 and PKS2 may be more similar between these species.

Polypropionates are abundant in molluscs worldwide and have been found in the sacoglossan species *E. chlorotica*, *E. diomedea* and *P. ocellatus*. We now have expanded their presence to *E. timida*. In sacoglossans, polypropionates appear not only to bind fixed carbon from the chloroplast via *de novo* photosynthesis (Ireland & Scheuer, 1979), but also to act via oxidative and photocyclization pathways (Ireland & Scheuer, 1979; Zuidema & Jones, 2005; Zuidema et al., 2005; Zuidema & Jones, 2006). These reactions may protect sacoglossans from damage caused by photosynthetic reactive oxygen products and may therefore play an important role in life with functional kleptoplasty (Zuidema & Jones; Zuidema et al., 2005; Zuidema & Jones, 2006; de Vries et al., 2015; Powell et al., 2018). Interestingly, Torres et al. (2020) found the mRNA of PKSs in the transcriptome of *E. timida* (de Vries et al., 2015; Rauch et al., 2017). However, transcriptomes only show the genes that are expressed in a given tissue at a given time, resulting in a subset of all genes present in the genome. After the polypropionate scaffold is produced, it is decorated by tailoring enzymes. A C-methyltransferase and cytochrome P450 are probably required to produce the large number of polypropionates in Sacoglossa (Li et al., 2023). As in other eukaryotes, the genes for the enzymes involved in the production of natural products are not adjacent to each other. This makes it difficult to identify the corresponding genes for the decorating tailoring enzymes (Torres et al., 2020). The genomic environments of the genes encoding PKS1 and PKS2 were searched using BLASTp, mainly yielding uncharacterised and hypothetical proteins with no indication of their catalytic activity. Only highly complete and continuous genome assemblies, such as that of *E. timida*, can provide a comprehensive picture of the genes present.

In the future, the *E. timida* genome assembly may help to shed light not only on polypropionates and their role in the functional kleptoplasty, but also on immune genes. Immune genes are well studied in cnidarians, and have recently been discussed in Sacoglossa, as the innate immune system probably plays an important role in the establishment of the process of photosymbiosis (e.g. Koike et al., 2004; Davy et al., 2012; Fransolet et al., 2012; Lehnert et al., 2014; van der Burg et al., 2016; Melo Clavijo et al., 2018; Melo Clavijo et al., 2020). Despite the fact that we are gaining more and more knowledge about the process of functional kleptoplasty, it is still unknown how sacoglossans precisely identify the chloroplast of *A. acetabulum* as their symbiont and not as a pathogen nor how chloroplasts are absorbed. Melo Clavijo et al. (2020) found that sacoglossans - including *E. timida* - have a divergent collection of specific scavenger receptors and the thrombospondin-type-1 repeat protein superfamily, comparable to photosymbiotic cnidarians (e.g. Miller et al., 2007; Wood-Charlon & Weis, 2009; Zou et al., 2009; Neubauer et al., 2016; Brennan et al., 2017; Brennan & Gilmore, 2018; Dimos et al., 2019). Furthermore, they detected species-specific candidate genes that may be important for the symbiont identification in sacoglossans. We investigated the presence of polypropionate encoding genes in *E. timida* and hope that our genome assembly can serve as a reference genome for immune gene studies in Sacoglossa, too. We expect the genome assembly to contribute to future genetic studies on kleptoplasty and to serve as a high-quality resource for studies on sacoglossans and molluscs in general.

## Acknowledgements

The present study is a result of the LOEWE Centre for Translational Biodiversity Genomics (LOEWE-TBG) and was supported through the program ‘LOEWE-Landes-Offensive zur Entwicklung Wissenschaftlich-ökonomischer Exzellenz’ of Hesse’s Ministry of Higher Education, Research, and the Arts (HMWK). We thank the Genome Technology Center (RGTC) at Radboudumc for the use of the Sequencing Core Facility (Nijmegen, The Netherlands), which provided the PacBio SMRT sequencing service on the Sequel IIe platform. We would also like to thank the Bioscientia Institut für med. Diagnostik GmbH, especially Prof. Dr. Hanno Jörn Bolz and Dr. Christian Betz, for providing the PacBio SMRT sequencing service on the PacBio Revio platform. Dr. Bruno Hüttel provided valuable advice and support in establishing the PacBio ultra-low input library protocol in our laboratory at the LOEWE-TBG Centre. We would especially like to thank Dr. Vesa Havurinne and Dr. Sónia Cruz for their support in establishing an *Acetabularia acetabulum* culture in our laboratory in Frankfurt. The funding code for LOEWE-TBG is: LOEWE/1/10/519/03/03.001(0014)/52.

## Supplements

**Supplemental Table S1.**
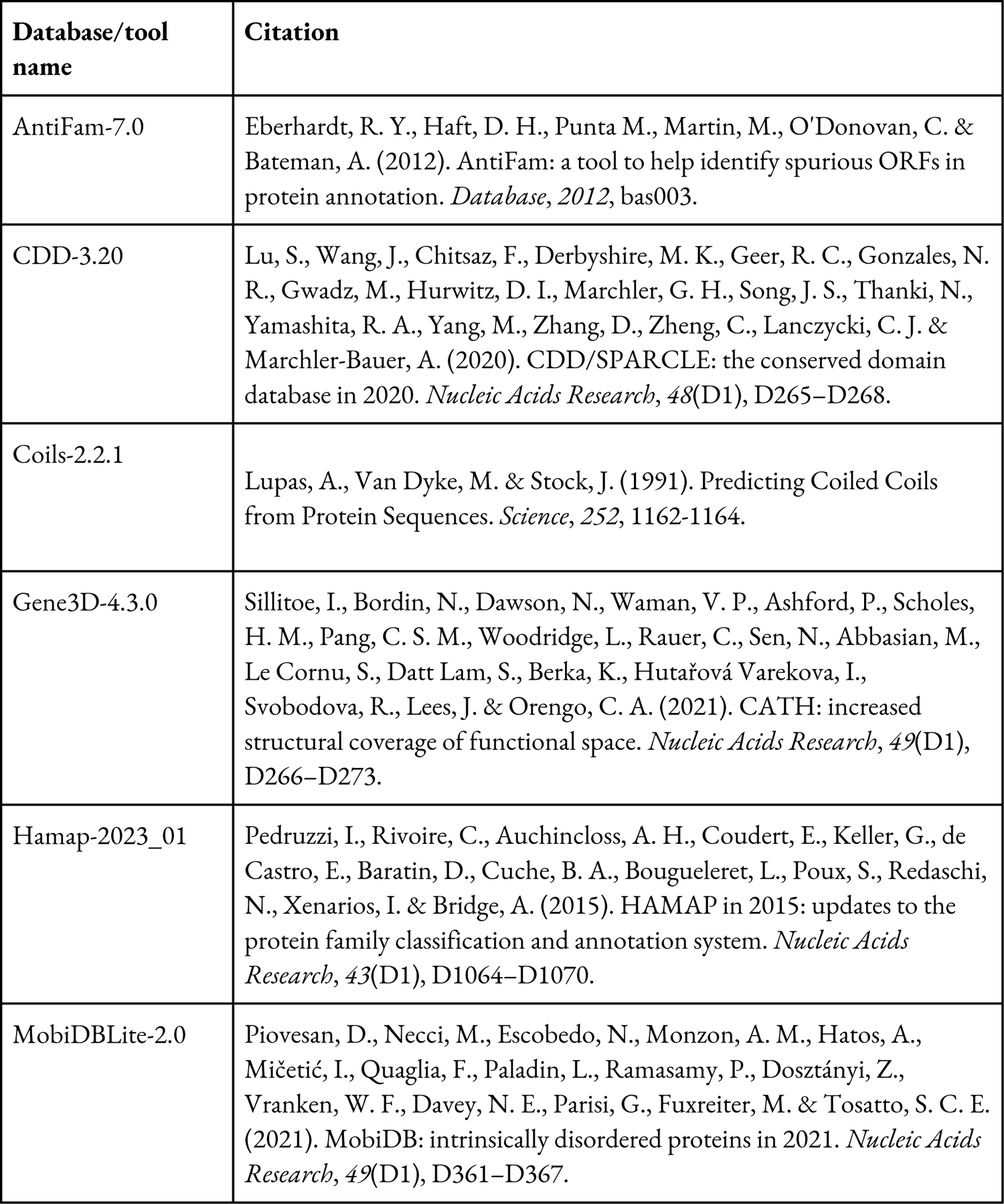
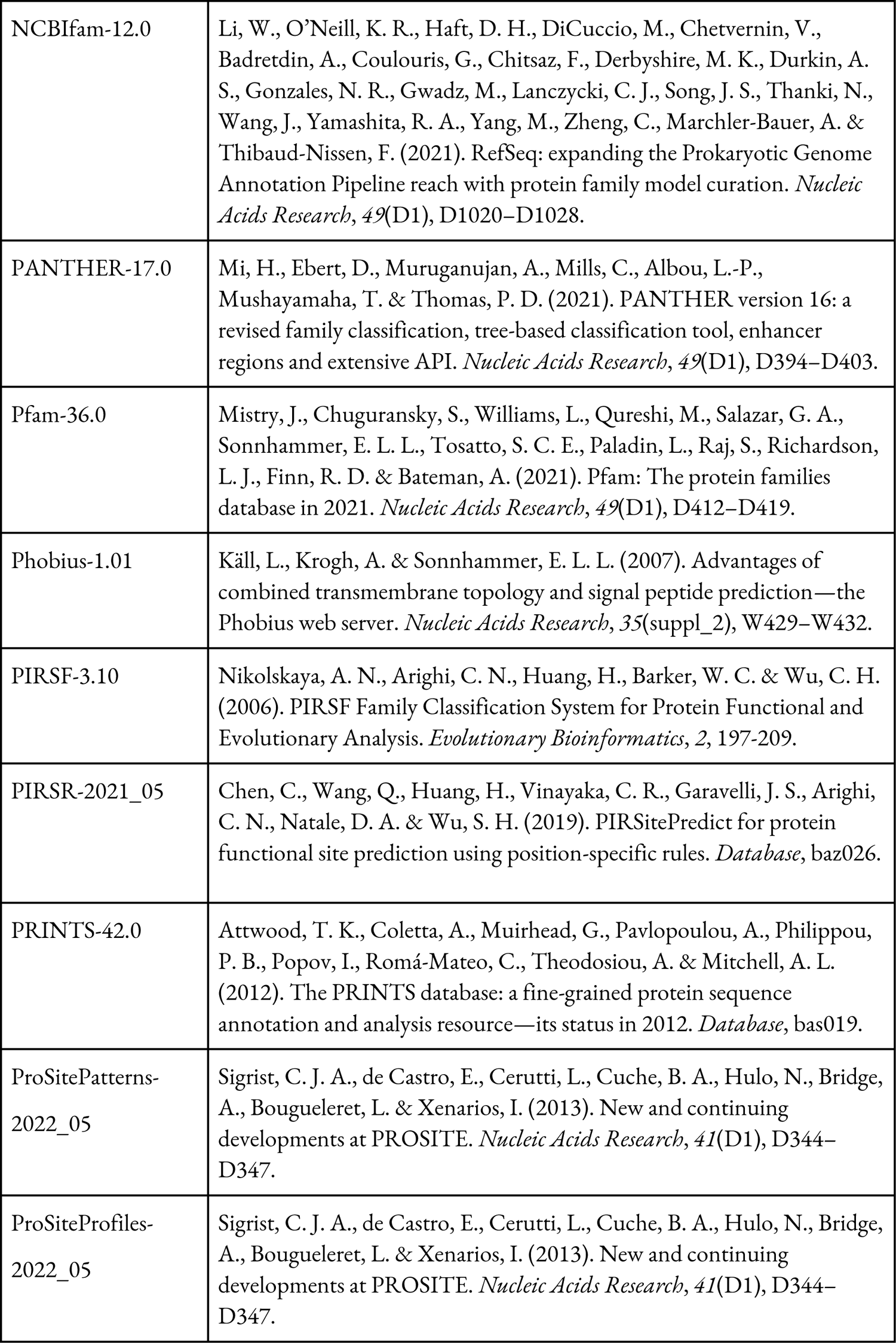
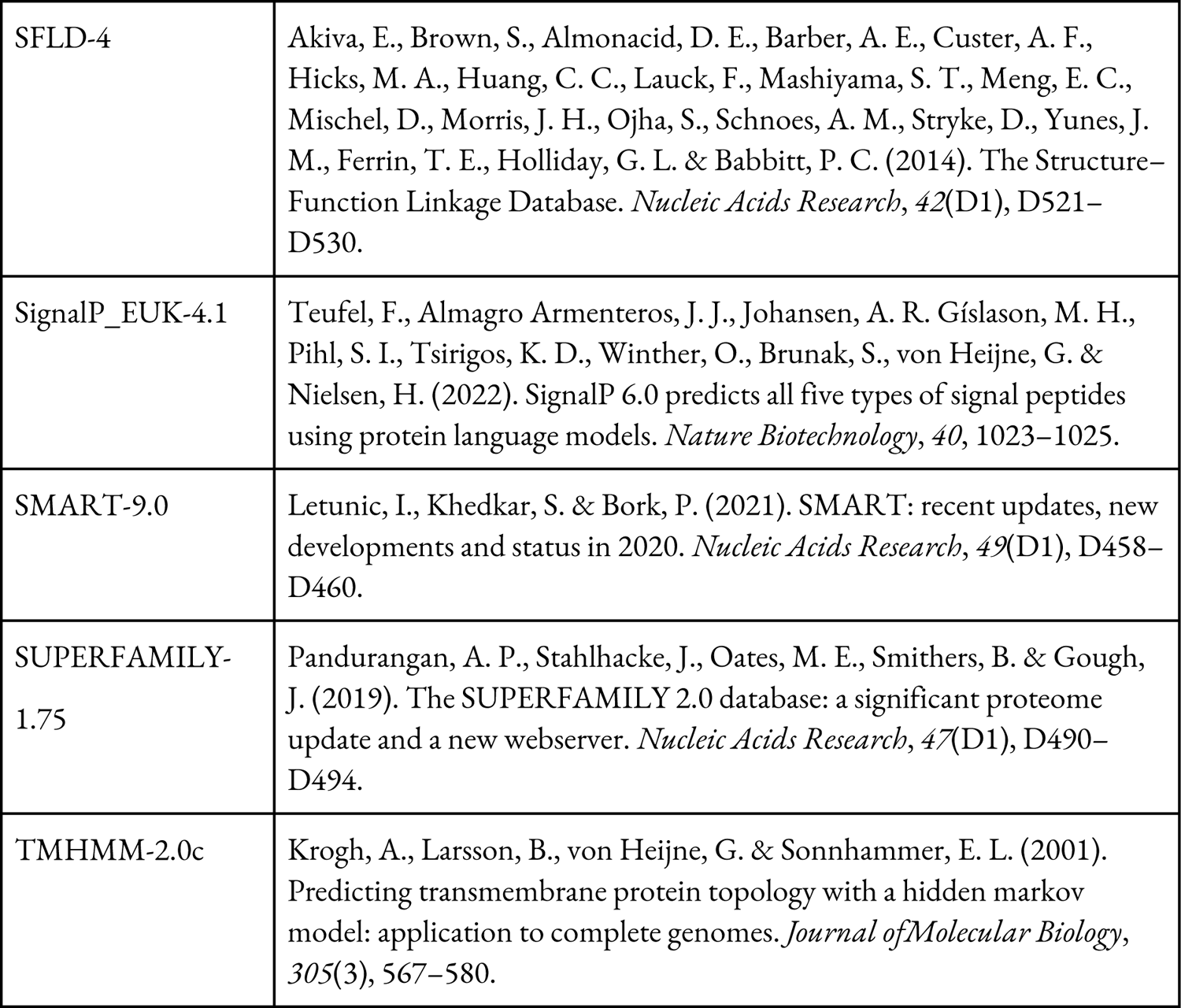
Databases and tools which were used while operating InterProScan version 5.64-96.0 (Jones et al., 2014).

**Supplemental Table S2.**
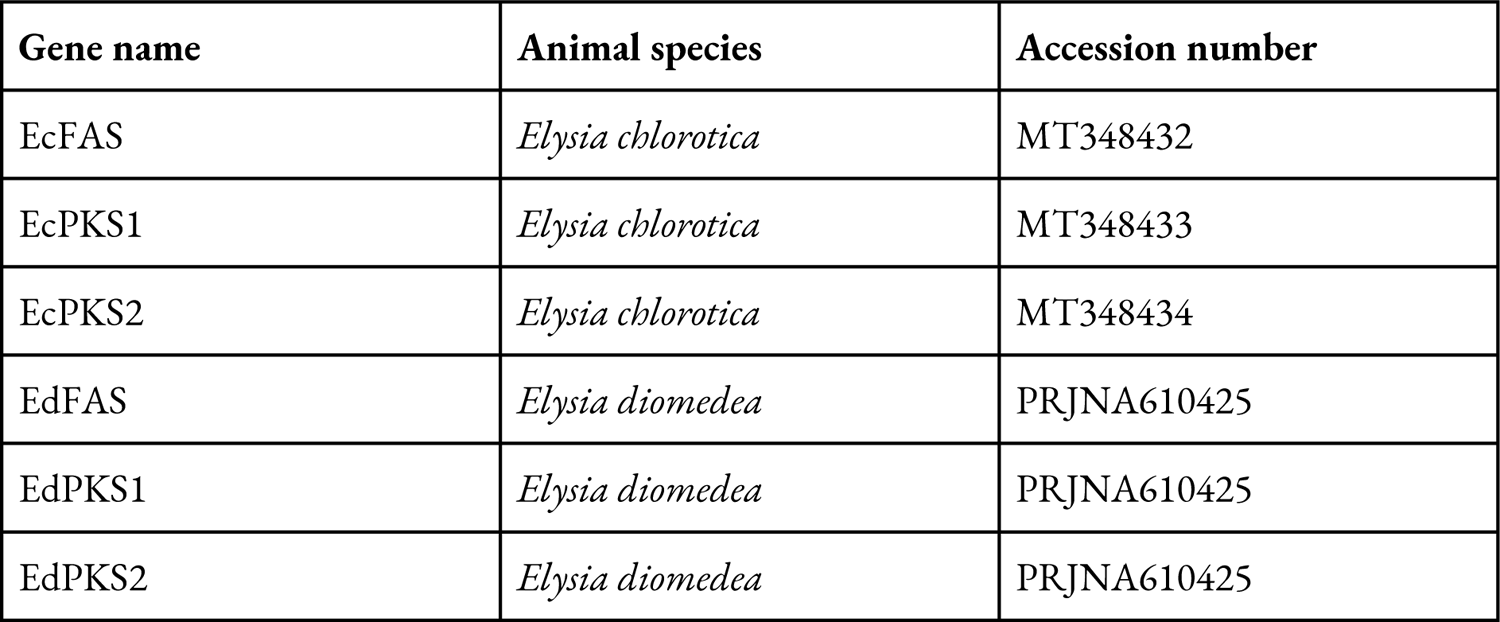
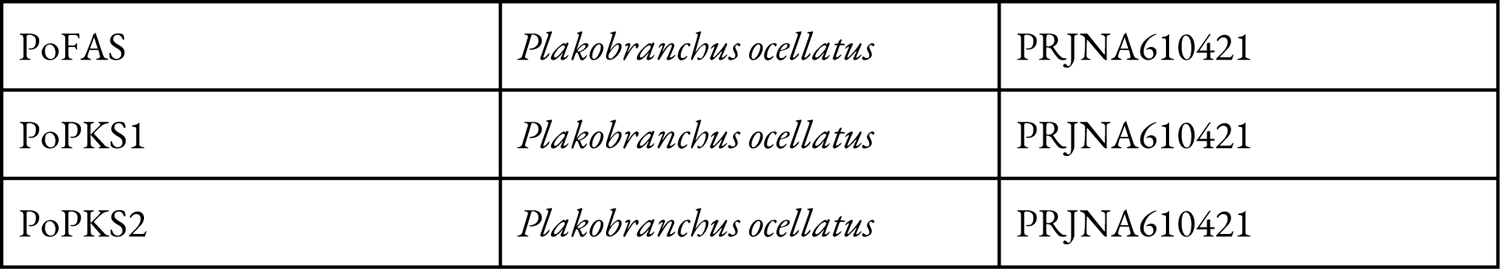
Table of PKS and fatty acid synthase (FAS) sequences from Torres, et al. (2020) including the animal species they were received from and the accession number.

**Supplemental Table S3.** Genome size estimates from two individuals of E. timida. The measured individual is given in brackets. Chopping buffer was prepared as described by Galbraith, et al. (1983). Propidium iodide was used as a fluorescent dye. We used the house cricket *A. domesticus* as standard reference (genome size: 2000 Mb).

| Organism (individual 1/2) | Genome size sample [Mb] | rCV value | Average genome size [Mb] |  |
| --- | --- | --- | --- | --- |
| *E. timida* (1) | 897.13 | 3.37% | 898.00 | 894.89 |
| *E. timida* (1) | 848.45 | 4.37% |  |  |
| *E. timida* (1) | 979.01 | 3.52% |  |  |
| *E. timida* (1) | 867.40 | 3.85% |  |  |
| *E. timida* (2) | 923.94 | 3.67% | 891.78 |  |
| *E. timida* (2) | 860.77 | 3.29% |  |  |
| *E. timida* (2) | 888.05 | 3.37% |  |  |
| *E. timida* (2) | 894.35 | 3.24% |  |  |

**Supplemental Table S4.** Sacoglossan heterozygosity values. The heterozygosity values from all species except for *E. timida*, were inferred by Theisen & Jensen (1991).

| Species | Heterozygosity [%] |
| --- | --- |
| *Elysia timida* | 0.794 |
| *Alderia modesta* | 0.23 |
| *Calliopaea oophaga* | 0.30 |
| *Elysia viridis* | 0.18 |
| *Ercolania nigra* | 0.42 |
| *Limapontia capitata* | 0.25 |
| *Limapontia depressa* | 0.20 |

**Supplemental Table S5.**
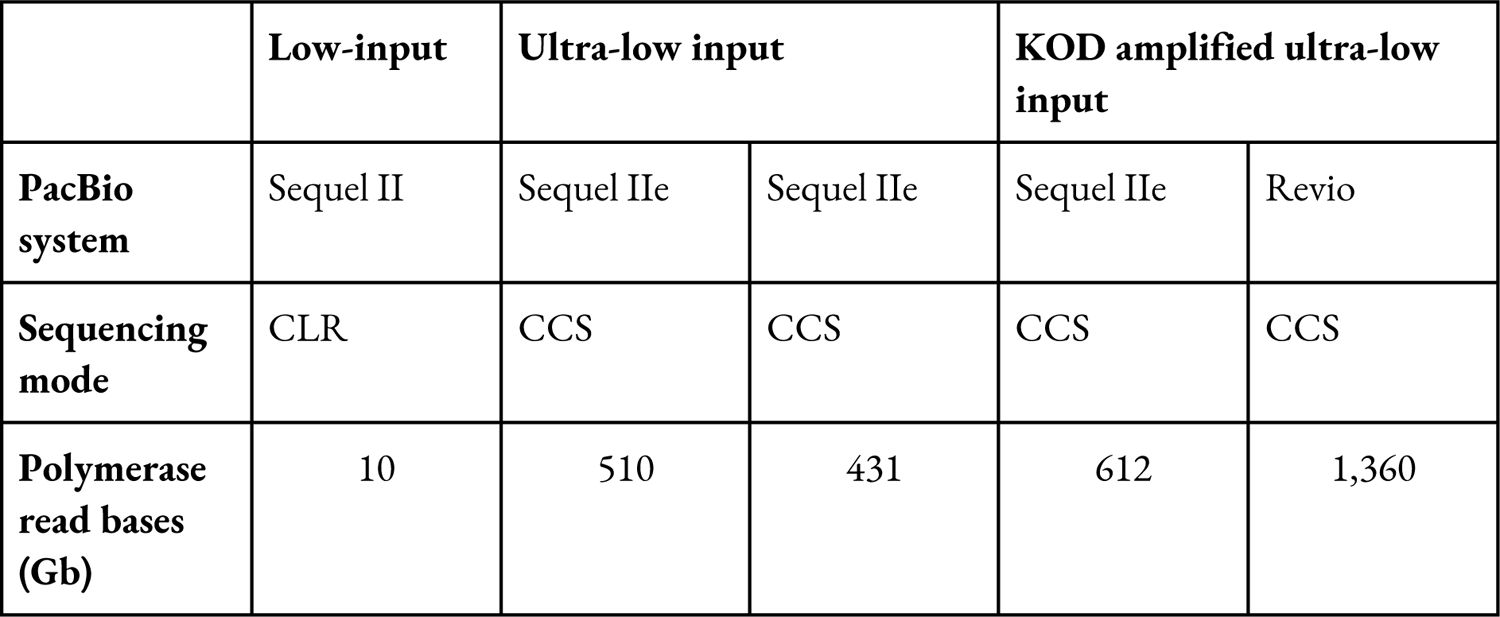
Sequencing output and subread mean length of the PacBio low- and ultra-input libraries.

|  | Low-input | Ultra-low input |  | KOD amplified ultra-low input |  |
| --- | --- | --- | --- | --- | --- |
| PacBio system | Sequel II | Sequel IIe | Sequel IIe | Sequel IIe | Revio |
| Sequencing mode | CLR | CCS | CCS | CCS | CCS |
| Polymerase read bases (Gb) | 10 | 510 | 431 | 612 | 1,360 |

**Supplemental Table S6.**
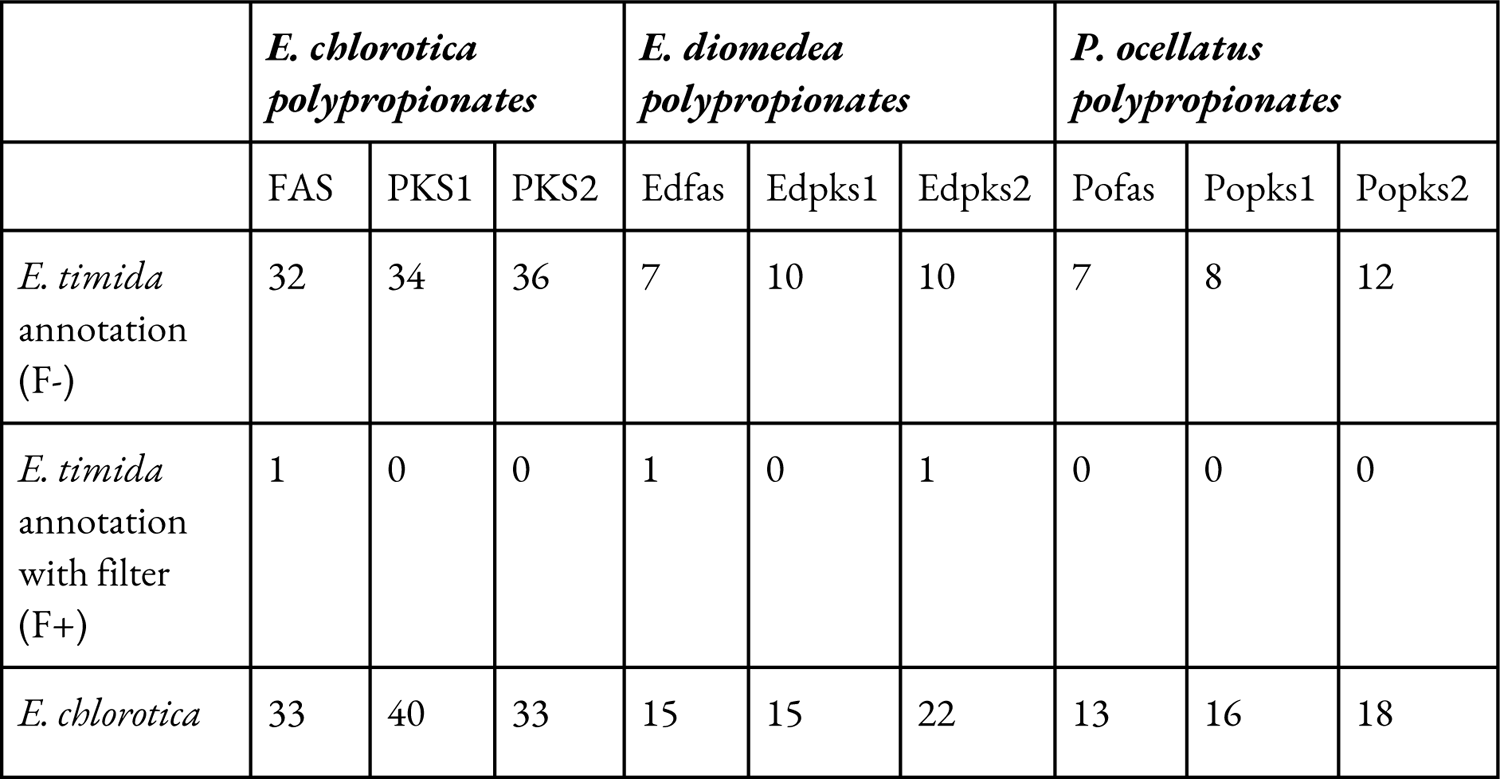
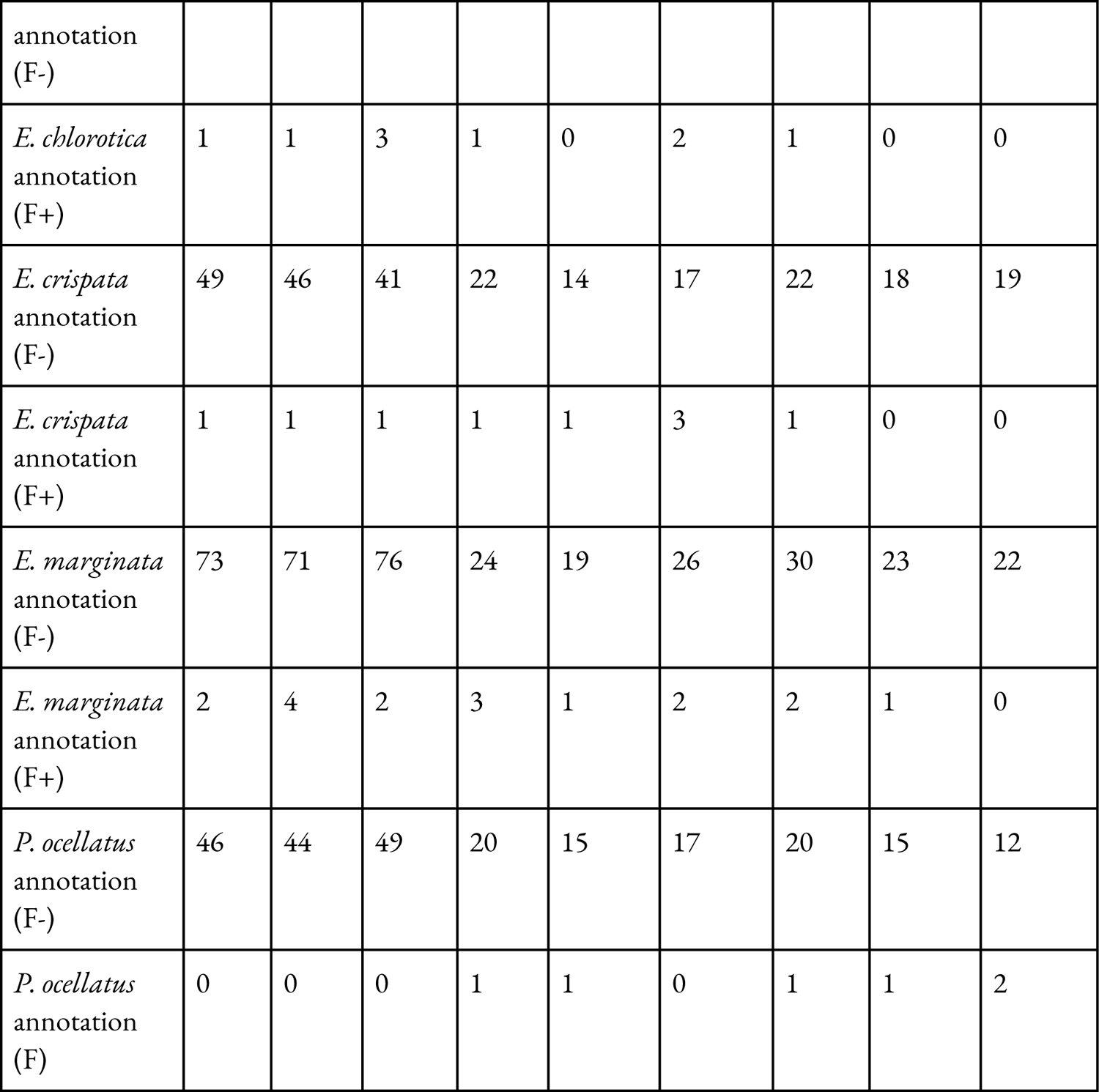
Result of the polypropionate blast search in the annotations of *E. timida*, *E. chlorotic*a, *E. diomedea* and *P. ocellatus*. Both unfiltered (F-) as well as filtered (F+) blast hits are shown.

**Supplemental Figure SX.**
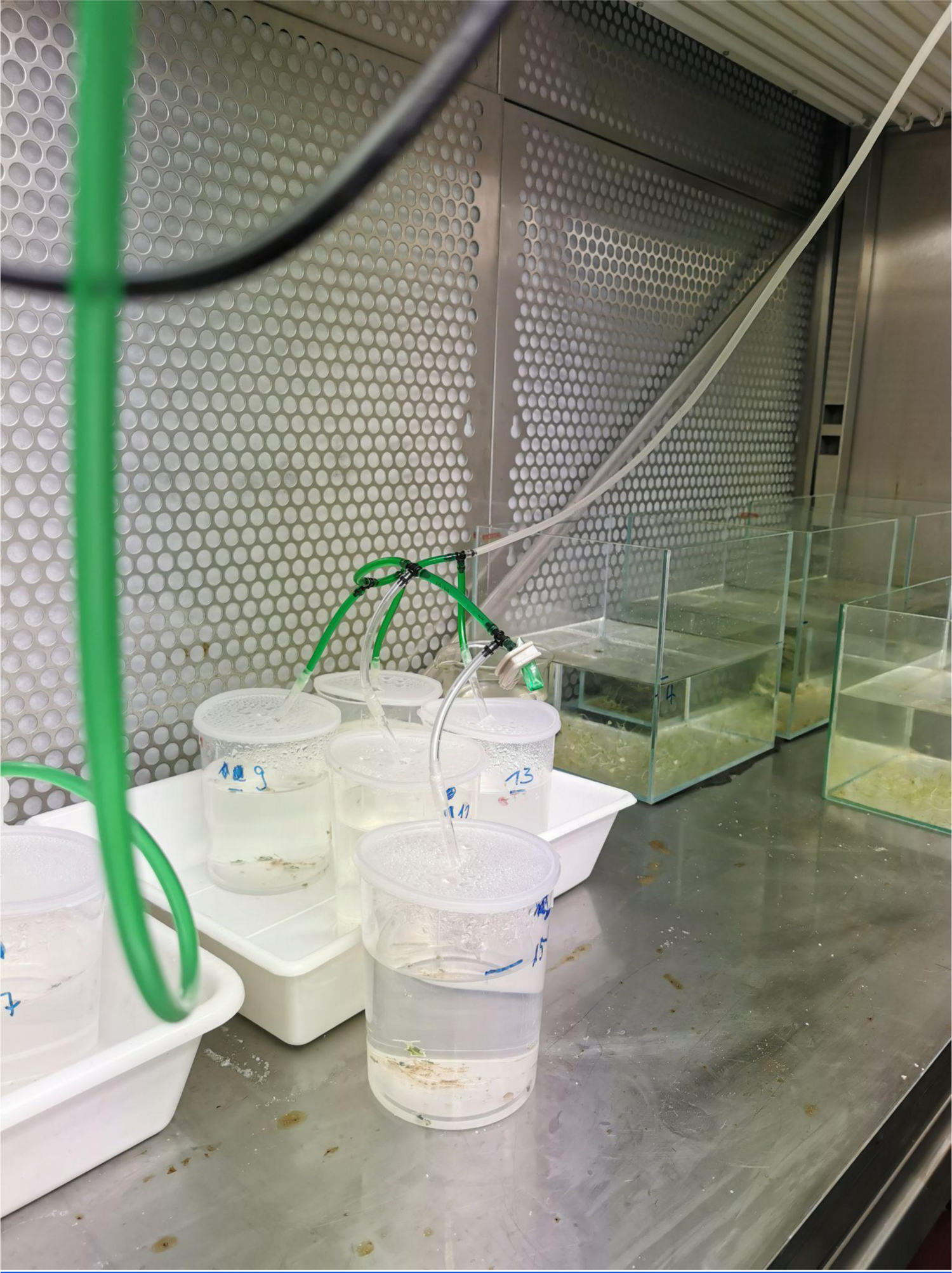
Climate chamber in which the *E. timida* slugs were kept in artificial sea water in plastic cups as aquariums. The green tubes provided the air supply.

**Supplemental Figure S2.**
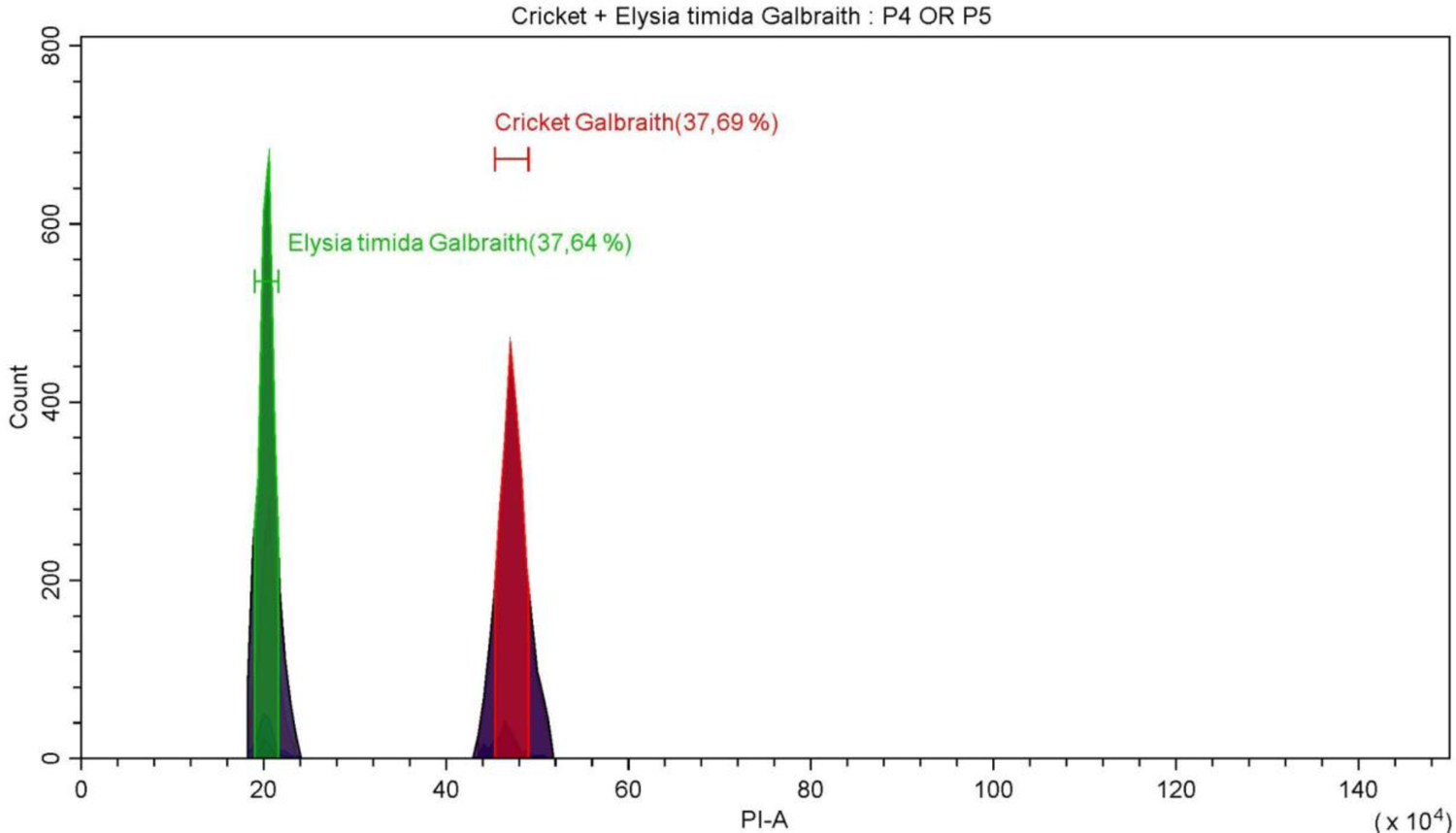
Genome size estimation of *E*. *timida* using flow cytometry. The histogram shows the relative propidium iodide fluorescence intensity obtained after simultaneous analysis of *E*. *timida* 2C (in green) and the house cricket *A. domesticus* 2C as an internal standard reference (in red). The PI fluorescent dyes were excited with a solid-state laser emitting at 488 nm. The y-axis gives the counts of propidium iodide (PI) stained nuclei. The x-axis displays the relative red PI fluorescence signal. To obtain the mean relative red PI fluorescence signals, the peaks were enclosed by line segments. The percentages in brackets are the portions of all events in the histogram enclosed by the respective line segments.

**Supplemental Figure S3.**
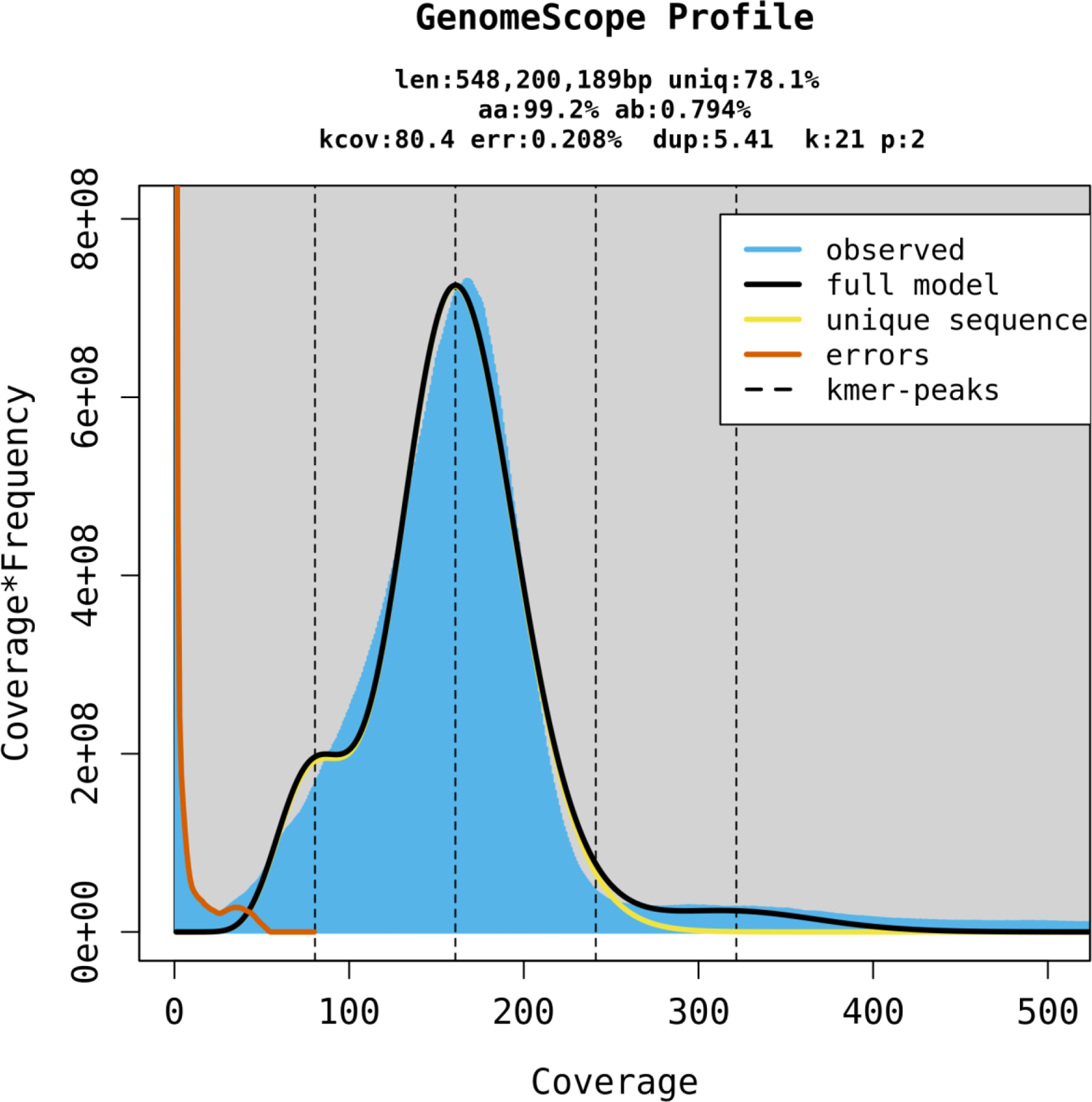
K-mer profile and estimates based on HiFi reads.

**Supplemental Figure S4.**
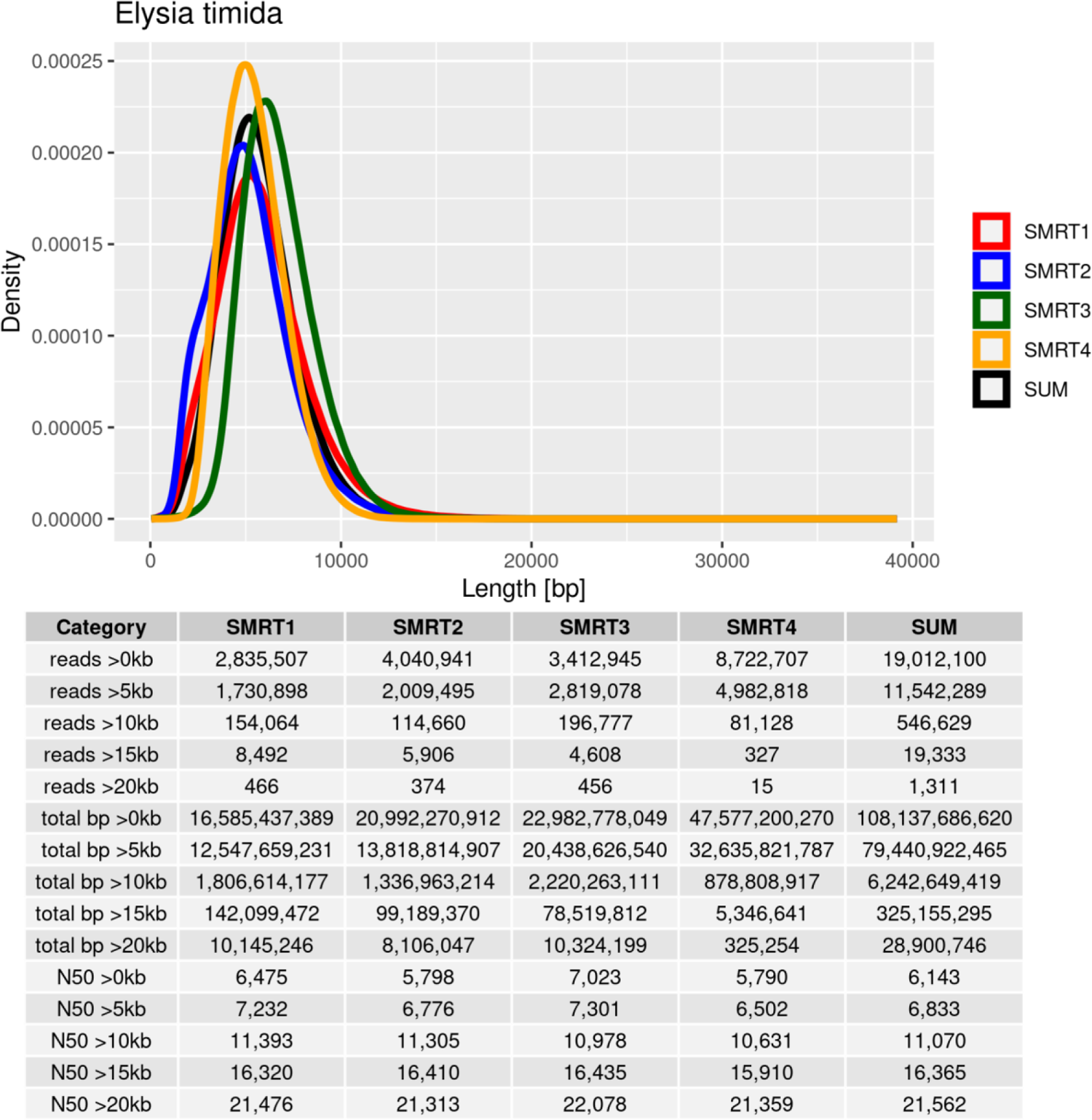
HiFi read length distribution and statistics. Standard PacBio ultra-low input libraries are listed as SMRT1 and SMRT2. PacBio ultra-low libraries amplified with KOD polymerase are shown as SMRT3 (Sequel IIe) and SMRT4 (Revio).

**Supplemental Figure S5.**
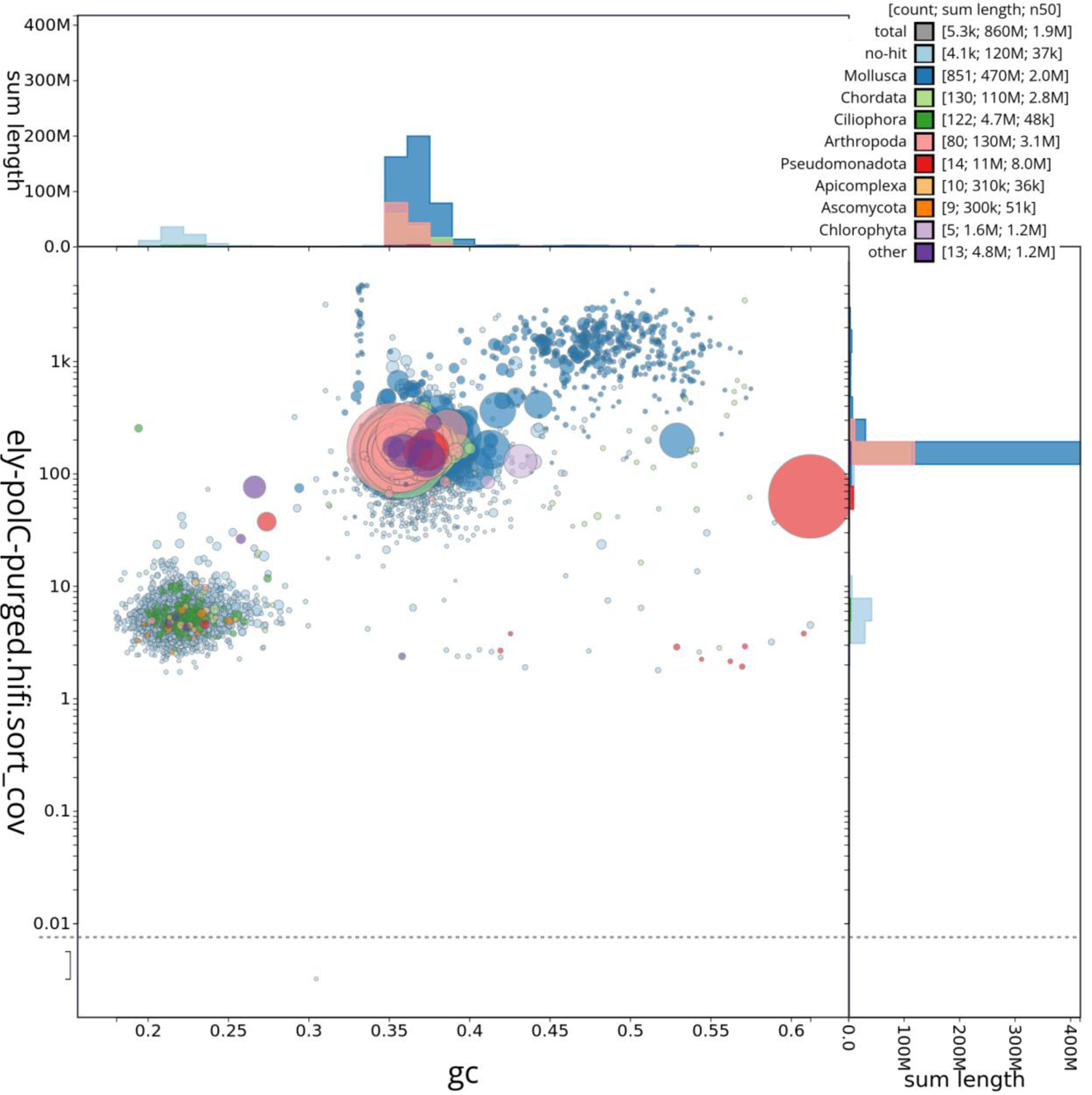
Blobplot of the assembly after polishing and purging. At this stage of the assembly process, contamination filtering with FCS was already conducted.

**Supplemental Figure S6.**
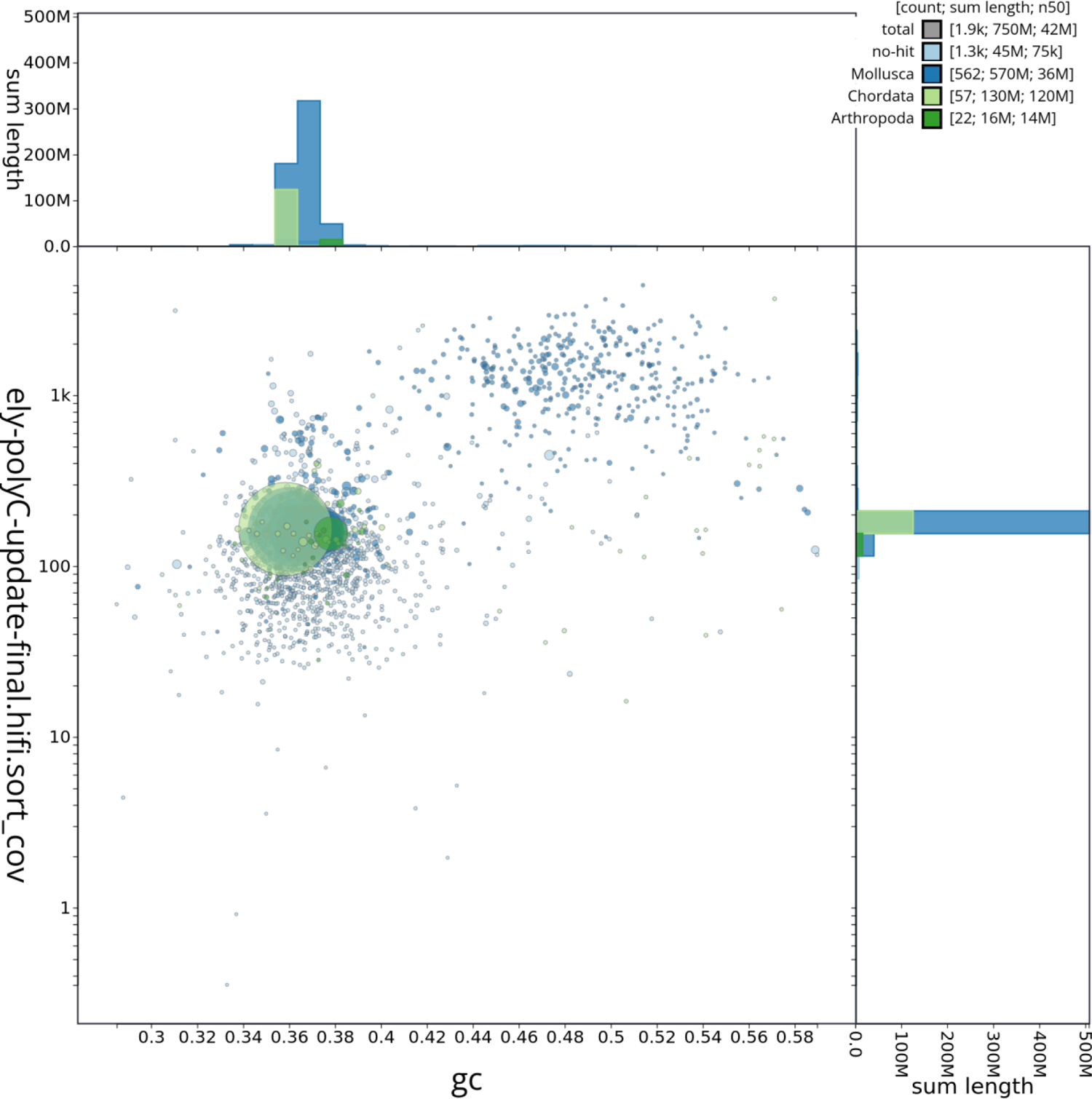
Blobplot of the final genome assembly.

**Supplemental Figure S7.**
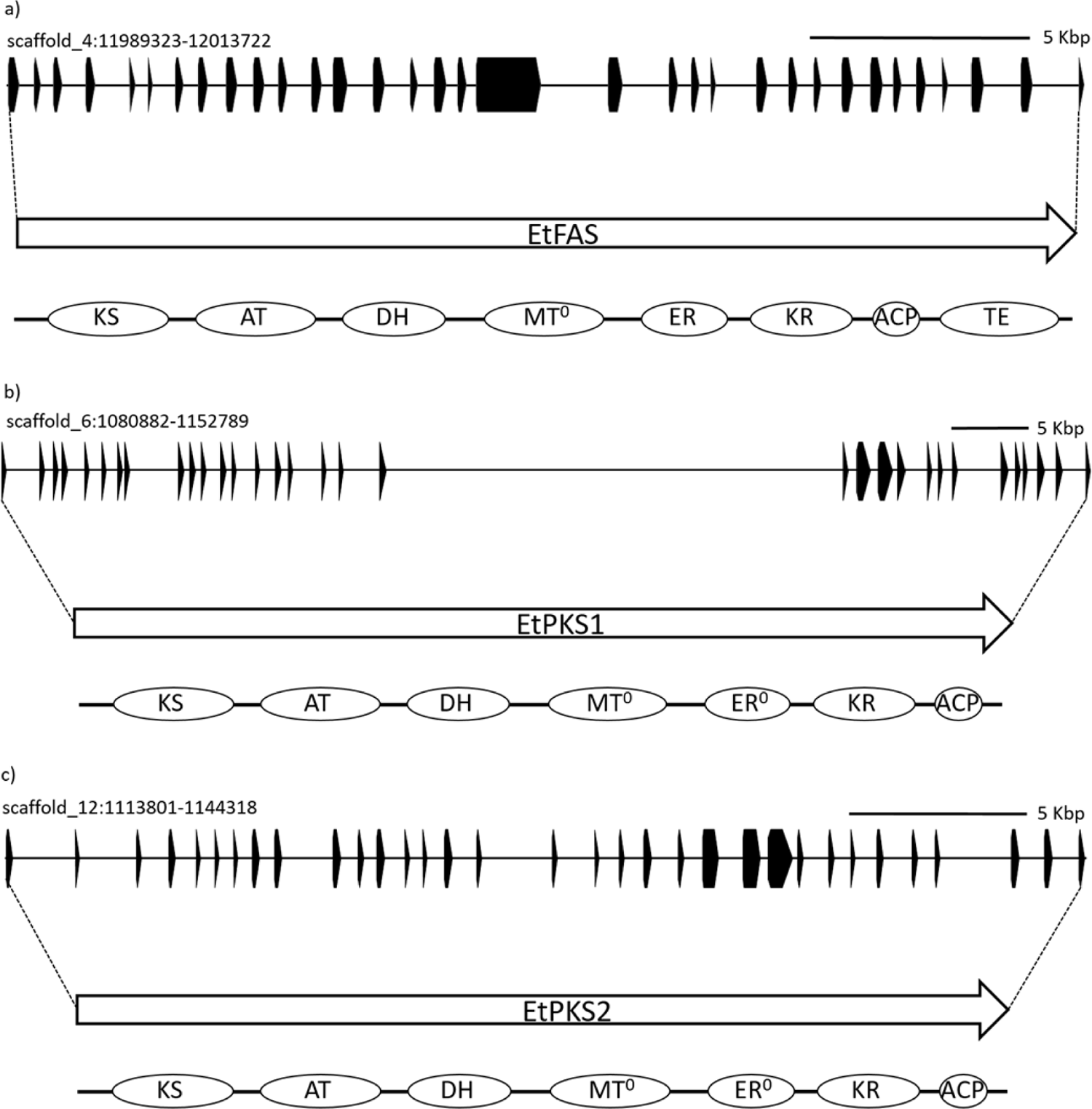
The genes encoding EtFAS, EtPKS1 and EtPKS2 are annotated in the genome of *E. timida*. The exons are labelled in black on the excerpt of the genomic sequence. The arrows present the transcript of the a) EtFAS, b) EtPKS1 and c) EtPKS2. The domains of the enzymes are presented in bubbles below the arrow. The gene encoding EtPKS1 was annotated manually based on sequence homology with the EcPKS1. The label with an x^0^ indicates an inactive domain.

**Supplemental Figure S8.**
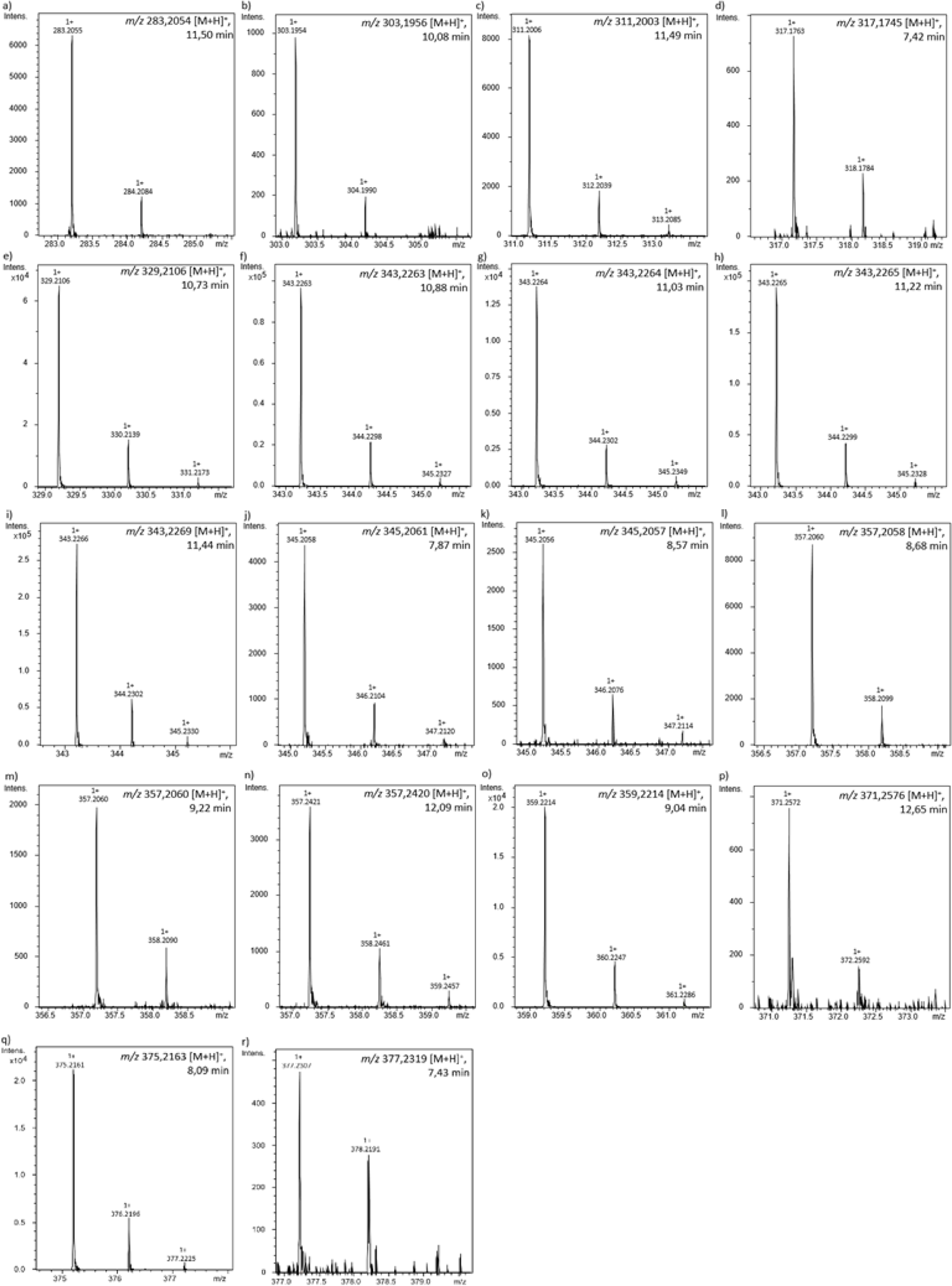
Isotopic patterns of the putative polypropionates produced by *E. timida* and identified by HPLC-ESI-HRMS analysis.

**Supplemental Figure S9.**
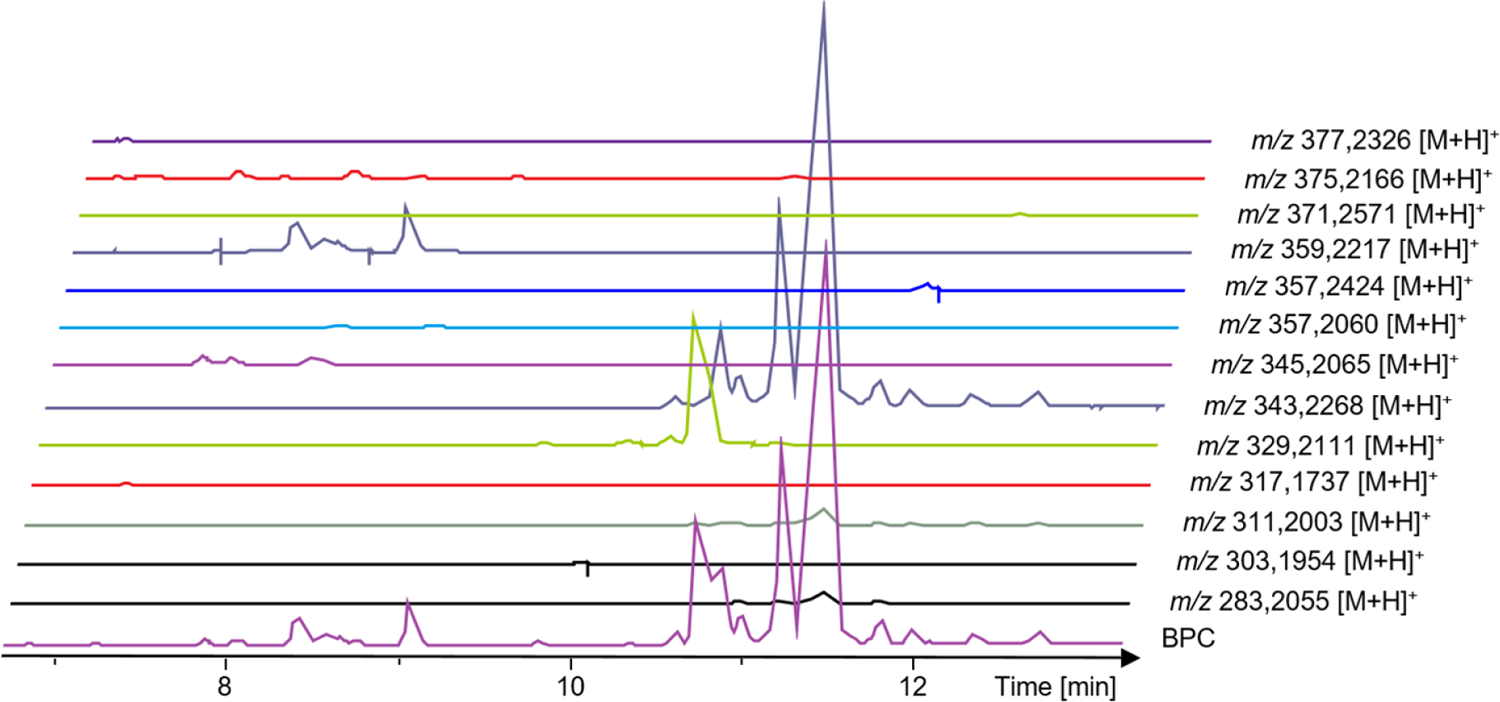
HPLC-MS data of *E. timida* extracts. BPC and EIC of polypropionates shown in figure (GNPS cluster).

**Supplemental Figure S10.**
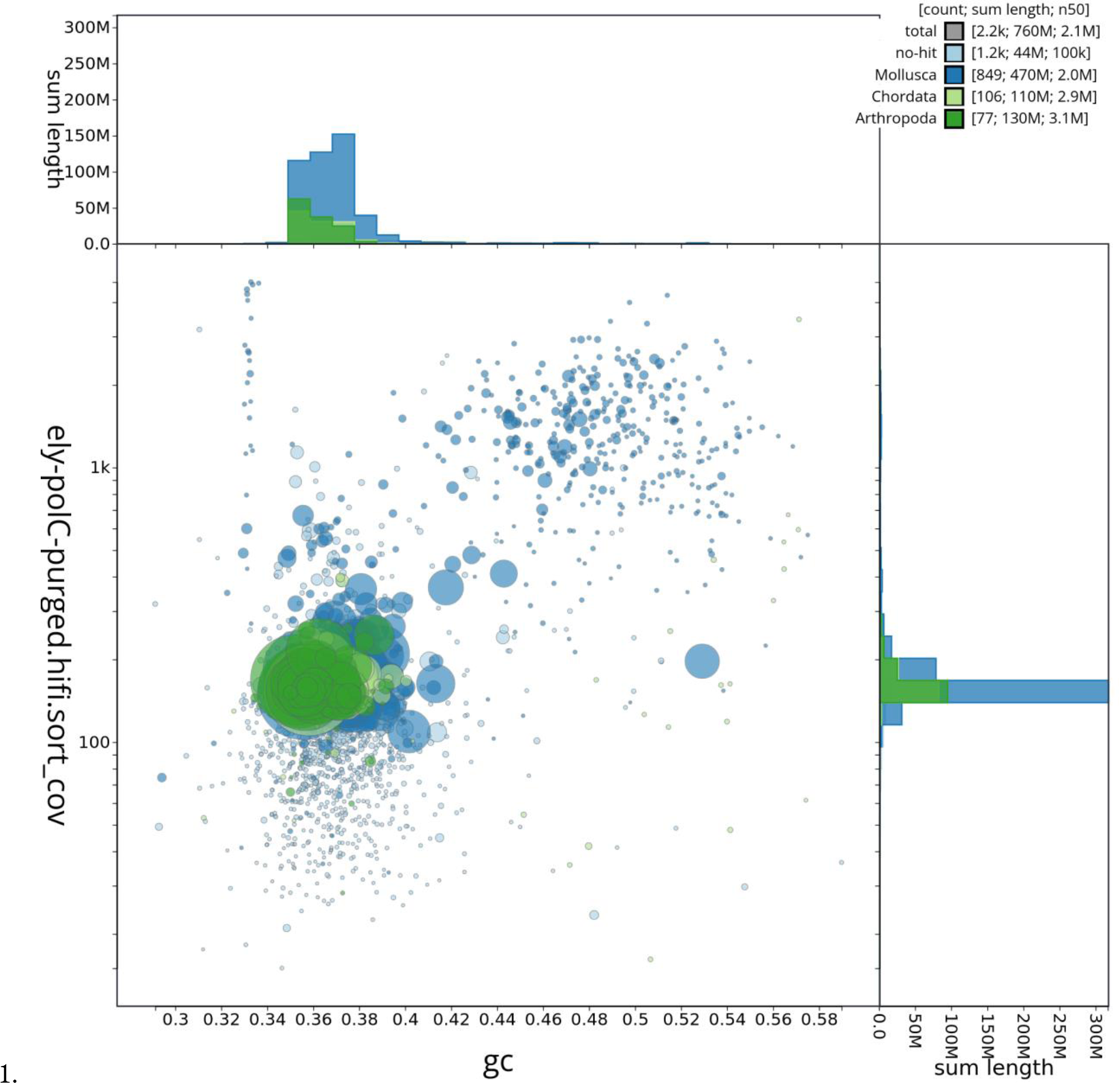
Blobplot of the assembly after removing sequences identified as contamination and before HiC scaffolding.

## Notes

### Competing Interest Statement

The authors have declared no competing interest.

